# Finding the perfect promoter: Correlating single-cell transcriptome data with gene drive performance

**DOI:** 10.1101/2025.10.08.681126

**Authors:** Yingke Wu, Yunchen Xia, Ziyin Yao, Weizhe Chen, Xihua Jia, Nan Liang, Jackson Champer

## Abstract

Gene drive can control pathogen transmission or suppress vector populations by spreading drive alleles with super-Mendelian inheritance. CRISPR homing drive currently represents the most powerful type, and regulating Cas9 expression with specific promoters has been effective for improving drive performance. However, selecting these is often a major challenge. Here, we evaluated 35 Cas9 constructs driven by distinct promoters in different gene drive systems and identified associations between drive performance and single-cell RNA expression patterns of the promoter-associated genes. Our results indicate that higher drive conversion is significantly associated with elevated expression of the promoter-associated gene in the respective reproductive cells, but embryo resistance allele formation correlates with excessive female germline expression. For males, early germline expression produces superior performance. Thus, we find that optimal drive performance requires restricting Cas9 expression to a tight quantitative and spatiotemporal window. In addition, found that *in situ* integrated *rhino*-Cas9 constructs significantly reduce somatic expression, underscoring the importance of genomic locus. On the basis of these results, we propose criteria for selecting promoters, providing a theoretical rationale and practical guidance for optimization of promoter elements in homing gene drive systems.

## Introduction

Gene drive is a genetic manipulation technology that leverages the characteristics of selfish genetic elements and modern gene-editing tools. It enables control of pathogens or suppression of vector populations by promoting the spread of drive alleles though populations by super-Mendelian inheritance^1–5^. Among various drive approaches, CRISPR based homing gene drives are the most extensively studied and powerful. In this system, Cas9 and one or more guide RNAs (gRNAs) cleave the wild-type allele in germline cells of heterozygotes, converted them into drive alleles and thereby increasing the drive inheritance probability beyond Mendelian expectations (though recent studies have also showed that others nucleases such as Cas12a can function efficiently ^6^). The feasibility of CRISPR based homing gene drive has been demonstrated in a variety of model organisms, disease vectors, invasive species, and agricultural pests including *Drosophila melanogaster*^6–9^, *Drosophila suzukii*^10^, *Ceratitis capitata*^11^, *Saccharomyces cerevisiae*^12,13^, *Candida albicans*^14^, *Plutella xylostella*^15^, *Anopheles gambiae*^16–18^, *Anopheles stephensi*^19,20^, *Aedes aegypti*^21^, *Culex quinquefasciatus*^22^, *Mus musculus*^23^, and viral populations^24,25^. However, none of these systems have yet been deployed in the wild. This is in large part due to issues in overall drive effectiveness. Developing drives that function well even in laboratory cage populations has been challenging, at least in multicellular organisms.

Gene drive performance can be evaluated from three aspects^26^: the drive conversion rate, the formation of resistant alleles, and fitness costs. The drive conversion rate is a core parameter for assessing the propagation potential of drive element within a population. It primarily reflects the efficiency with which Cas9 cleaves the target DNA in germline cells and precisely inserts the drive element through homology-directed repair (HDR). A high conversion rate enhances deviation from Mendelian inheritance ratios, thereby accelerating the spread of the drive element throughout the population^5^. This can allow a modification drive to spread despite fitness costs. For suppression drives, it is even more critical, directly affecting suppressive power.

Resistance alleles arise from the end-joining repair pathway or (more rarely) incomplete HDR, changing the wild-type sequence and thereby rendering them resistant to CRISPR-mediated cleavage, which interferes with drive spread^27,28^. Resistance alleles are classified into functional and nonfunctional resistance alleles^29,30^. Functional resistance alleles usually result from small indels at or near the Cas9 cut site that mutate the PAM or gRNA seed region, preventing cleavage while largely preserving the encoded protein. Such alleles can cause major drive failure, but they can be effectively overcome by gRNA multiplexing, though this can come at the cost of a modest decline in drive efficiency is many gRNAs are required^29,31^. By contrast, more common nonfunctional alleles remain problematic. They arise from frameshifts or more impactful indels that destroy protein function. All these alleles primarily form during the germline or later during embryonic stages^29,32^. In the germline, Cas9 can convert wild-type alleles into germline resistance alleles following DNA cleavage, enabling these resistance alleles to be heritable within the population^7^. During embryonic development, maternally deposited Cas9 and gRNA can mediate DNA cleavage and repair in the zygote or early embryo, resulting in the formation of embryo resistance alleles^19^. Resistance alleles formed during the zygote stage affects all cells in the developing organism, but it is also possible for resistance alleles to form later in all or nearly all cells. In both these cases, they can be part of the germline, potentially blocking drive conversion while still being inherited by the next generation. However, some only arise within somatic cells during embryogenesis and remain confined at the individual level, unable to enter the germline or be effectively transmitted to offspring^33^. These resistance alleles can still have similar negative fitness effects like somatic Cas9 activity (see below). Overall, high rates of resistance allele formation pose a major barrier to the successful spread of gene drives within populations, particularly when the drive system carries fitness cost^28,34^.

Fitness costs can come from a variety of sources, in particular somatic expression in suppression drives^29,35^. An ideal gene drive should ensure that Cas9 expression is restricted to germline cells associated with gametogenesis, but “leaky” somatic expression can occur even with a mostly germline-specific promoter^36^. This reduction in fitness reduces survival and/or reproductive competitiveness, ultimately affecting the efficiency of the drive^29,37^. Aberrant expression of Cas9 in somatic tissues may lead to reduced fitness if the wild-type allele (which is destroyed by somatic expression, regardless of whether HDR or end-joining occurs) is required for survival or fertility. Thus, this fitness cost due to somatic activity is particularly problematic for suppression gene drives^17,30,37^. Suppression drives generally rely on the rapid spread of the drive within the population to accumulate a sufficient proportion of sterile or nonviable individuals, thereby achieving effective population suppression (high “suppressive power”). However, if drive/wild-type heterozygous individuals carrying the drive suffer fitness cost due to somatic activity, the drive will spread to a lower proportion of the population when it reaches equilibrium, potentially preventing population elimination^38,39^. Additional fitness costs may result from the burden or toxicity of transgenes, Cas9 off-target effects, and insertion-site position effects^40^.

The Cas9 promoter plays a critical role in determining the drive performance^36^. To date, multiple Cas9 promoters with different drive performance have been identified in *Drosophila melanogaster*. For example, the *nanos* promoter exhibits a relatively high drive conversion rate and embryo resistance rate, while maintaining low levels of leaky somatic expression^7^. In contrast, the *CG4415* promoter achieves a similarly high drive conversion rate while effectively reducing the embryo resistance rate^36^. However, it also maintains low levels of somatic expression, higher than that observed with the *nanos* promoter. In *Anopheles gambiae*, the *nanos* and *zpg* promoters also have high drive conversion efficiency and low embryo resistance, while still retaining some somatic expression that causes fitness costs in suppression drives^37,41^. Although previous studies have revealed that different promoters exhibit distinct performance in gene drive systems^36^, the regulatory mechanisms by which Cas9 promoters influence drive efficiency at the transcriptomic level remain unclear, and systematic efforts to screen for high-performance promoters remains limited. This challenge is particularly evident in non-model species with limited foundational research^42–44^, where Cas9 promoters are typically sourced from those validated in model organisms or identified by cloning the upstream regions of highly expressed reproductive tissue genes using RNA-seq data. However, these approaches have disadvantages. On the one hand, the expression levels and spatiotemporal patterns of promoters transferred across species may not always align with expectations^20,36^. On the other hand, RNA-seq screening only reflects the overall average expression levels at the tissue level, failing to resolve the fine-scale expression patterns of genes across different cell types and developmental stages within complex reproductive tissues where critical germline cells represent only a fraction of the total cells that are assessed.

Single-cell transcriptome data can potentially address this issue, giving precise information on expression patterns. Yet, it remains unclear exactly what expression pattern may be optimal and what kinds of differences may be found between such data and Cas9 activity when driven by promoters from native genes. Therefore, it is crucial to further investigate the relationship between expression of candidate promoter genes and drive performance to develop efficient and broadly applicable Cas9 promoter screening strategies for diverse species.

Based on this, our study systematically tested 35 Cas9 expression constructs driven by different promoters in several different types of gene drive systems. By comparing their performance and integrating single-cell transcriptomic data^45^, we identified potential correlations between drive performance and gene expression patterns. While high expression in specific reproductive cells is necessary for efficient drive conversion, excessive or ectopic expression generates undesirable resistance or somatic activity. This leads to our central conclusion that optimal performance requires confining Cas9 expression to a tight quantitative and spatiotemporal window. Our analysis enables us to define the key criteria for selecting high performance promoters, providing a foundation and practical strategies for optimization of nuclease promoters in homing drives.

## Methods

### Plasmid construction

The plasmids SNc9CGnG and BHDcN1 were previously constructed^29,36^. They were first digested at unique sites with restriction enzymes (New England Biolabs). Genomic or plasmid DNA of interest were amplified using NEBNext^®^ Ultra™ II Q5^®^ Master Mix. Gibson assembly was performed using NEBuilder^®^ HiFi DNA Assembly Master Mix. The assembled plasmids were transformed into chemically competent *E. coli* strains DH5α (TIANGEN) or XL10 (Angyu). Plasmids were extracted using the ZymoPure Midiprep Kit and verified by Sanger sequencing or full-plasmid sequencing.

### Generation of transgenic lines

Embryo microinjections were performed by UniHuaii and Fungene Biotech. For Cas9 promoter lines, donor plasmids (500 ng/µL) were co-injected with the gRNA helper plasmid BHDabg1^46^ (100 ng/µL) and the Cas9 expression plasmid TTChsp70c9^47^ (450 ng/µL) into *w^1118^* embryos. For the RON line, donor plasmid ISc9ROG (500 ng/µL) were co-injected with the gRNA helper plasmid BHDrhg2 (100 ng/µL) and the Cas9 expression plasmid TTChsp70c9 (450 ng/µL) into *w^1118^* embryos. For the *cinnabar* gRNA line, the donor plasmid (640 ng/µL) was co-injected with TTChsp70c9 (460 ng/µL) into *w^1118^* embryos. Injected individuals were crossed to *w^1118^* flies, and progeny carrying the drive element were identified by fluorescence. After confirming the insertion site by Sanger sequencing of genomic DNA, homozygous lines were established by inbreeding for most lines. However, a homozygous RON line could not be established as the Cas9 insertion disrupts the *rhino* locus, which results in recessive female sterility^48,49^. The gRNA lines targeting *yellow*^7^, *doublesex*^50^, and *RpL35A*^8^ were previously generated. Previously constructed Cas9 promoter lines include: SNc9nG, SNc9SnG, SNc9XnGr, SNc9VnG, SNc9EnG, SNc9CnG, SNc9DnG^36^, and SNc9320nG. Plasmid maps of all new sequences are available on GitHub (https://github.com/jchamper/Single-Cell-Promoters-Homing).

### Fly rearing and phenotypes

Flies were reared on modified Cornell standard cornmeal medium (using 10 g agar instead of 8 g per liter, addition of 5 g soy flour, and without the phosphoric acid) and maintained in vials at 25 °C under a 14/10 h day/night cycle at 60% humidity. Flies were anesthetized using CO₂ and screened for fluorescence using NIGHTSEA adapter SFA-GR for DsRed and SFA-RB-GO for EGFP. Fluorescent protein expression was specifically driven in the white eyes of *w^1118^* under the control of the 3×P3 promoter and the SV40 3’UTR/terminator. EGFP expression indicates the presence of the Cas9 allele, while DsRed indicates the presence of the drive element in individual flies.

In the *yellow* system, loss of *yellow* (*y*) function yields a recessive yellow body phenotype characterized by a lack of dark pigment in adults. We scored body color to infer resistance genotypes and somatic activity. Because *y* is recessive and X-linked, hemizygous males with a disrupted *y* are yellow, whereas females are yellow only when both X-linked copies are disrupted. Resistance alleles were grouped as functional or nonfunctional, of which only the latter contributes to the yellow phenotype. In addition, post-zygotic cleavage and end-joining can occur in embryos, producing flies with partial loss of *y* function. Somatic activity can also result in cleavage of only some cells in the body. In both these situations, individuals were scored as “mosaic”. In practice, flies typically display either wild-type or yellow body and wing pigmentation, but some flies may exhibit a mosaic pattern with mixed yellow and wild-type pigmentation on the dorsal abdomen or wings, which were classified as “mosaic”. In the *cinnabar* system, individuals lacking wild-type alleles usually have a bright orange eye phenotype, but this is not present in *w^1118^* individuals which do not produce eye pigments.

### Drive performance assays

Males from different promoter lines were crossed with females from various gRNA lines to generate males or female virgins carrying both DsRed and EGFP fluorescent markers. In parallel, somatic expression in the *yellow* system was evaluated by examining the body color of female virgins carrying both fluorescent markers. These individuals were then crossed to the *w^1118^* strain, and their progeny were scored based on fluorescence markers and other relevant phenotypes. In the *doublesex* system, somatic activity was indirectly assessed by the fertility of female virgins carrying both fluorescent markers.

To control for batch effects, statistical analysis was performed in R (version 4.5.0) following established approaches^8,31^. A generalized linear mixed-effects model (GLMM) was fitted using the lme4 package. Each individual was classified into binary categories (e.g., drive vs. non-drive, resistance vs. non-resistance), with vial ID included as a random effect (drive ∼ 1 + (1 | group)). After model fitting, marginal means and their standard errors were estimated using the emmeans package to calculate the expected probabilities for each genetic outcome. The program used is available on Github (https://github.com/jchamper/Single-Cell-Promoters-Homing).

### Cell type-specific differential expression analysis

We obtained the single-cell expression matrix of the Fly Cell Atlas from the Single Cell ExpressionAtlas^45^ (https://ftp.ebi.ac.uk/pub/databases/microarray/data/atlas/sc_experiments/E-MTAB-10519/E-MTAB-10519.aggregated_filtered_normalised_counts.mtx). Cell type annotations were obtained from the associated metadata file (https://www.ebi.ac.uk/gxa/sc/experiment/E-MTAB-10519/download?fileType=experiment-design&accessKey=). Gene expression values were then aggregated by cell type by calculating the median expression level for each gene within each cell type. The bulk RNA-Seq data were obtained from FlyBase database (release fb_2024_01)^51,52^.

Promoter-associated genes were classified based on drive performance as follows. For female germline cut rate (drive conversion + germline resistance allele formation), genes with cut rates greater than 90% and less than 25% were compared. For female drive conversion, genes with conversion rates greater than 40% and less than 15% were compared. For male drive conversion, genes with conversion rates greater than 40% and less than 10% were compared. For embryo resistance, genes with drive conversion rates above 40% were further divided based on embryo resistance levels: greater than 65% versus less than 5%. For Cas9 somatic expression, genes with drive conversion rates above 40% were further divided based on body color: wild-type body color (zero/low somatic expression) versus fully yellow body color (high somatic expression). We deliberately contrasted extreme groups (very high and very low cut rate/drive conversion rates) to maximize effect size and improve statistical power. For the embryo resistance and somatic expression analyses, we included only promoters whose drive conversion rate met a predefined threshold, thereby excluding promoters that appeared to have low embryo resistance or somatic expression solely due to insufficient germline Cas9 expression that could not support acceptable drive efficiency and thus did not represent a good candidate for optimal drive performance.

For each of these pairs of groups, transcripts per million (TPM) differences between them were analyzed across various cell types. To mitigate the effect of zeros, raw TPM was transformed to log_2_(TPM + 1). For each cell type, expression values for the two gene groups were extracted, and differences were assessed for statistical significance using the Wilcoxon rank-sum test. In parallel, the effect size was defined as the between-group difference in this quantity, denoted Δlog _2_ (TPM+1). A cell type was considered to show significant differential expression if *p* < 0.05 and | Δlog_2_(TPM+1)| > 1.

## Results

### Selection of 35 promoter-candidate genes with different expression patterns

Based on single-cell transcriptomic data in the Fly Cell Atlas^45^, we selected 25 genes sex-specific or both-sex germline expression chosen to minimize somatic expression (Fig.1, Fig.S1). We included eight additional germline promoters from previous studies^36^ and added two constitutive genes (*Act5C* and *Ubi-p63E*) that are expressed at relatively high levels across various cell types and conditions. The promoter regions were defined as the 5′ untranslated region (5′ UTR) together with 64 to 3382 base pairs upstream of these genes (Table S1), selected according to transcription factor binding densities from the ReMap database (version 2022)^53^ while attempting to avoid regions in the coding sequence and UTRs of adjacent genes.

**Fig. 1:**
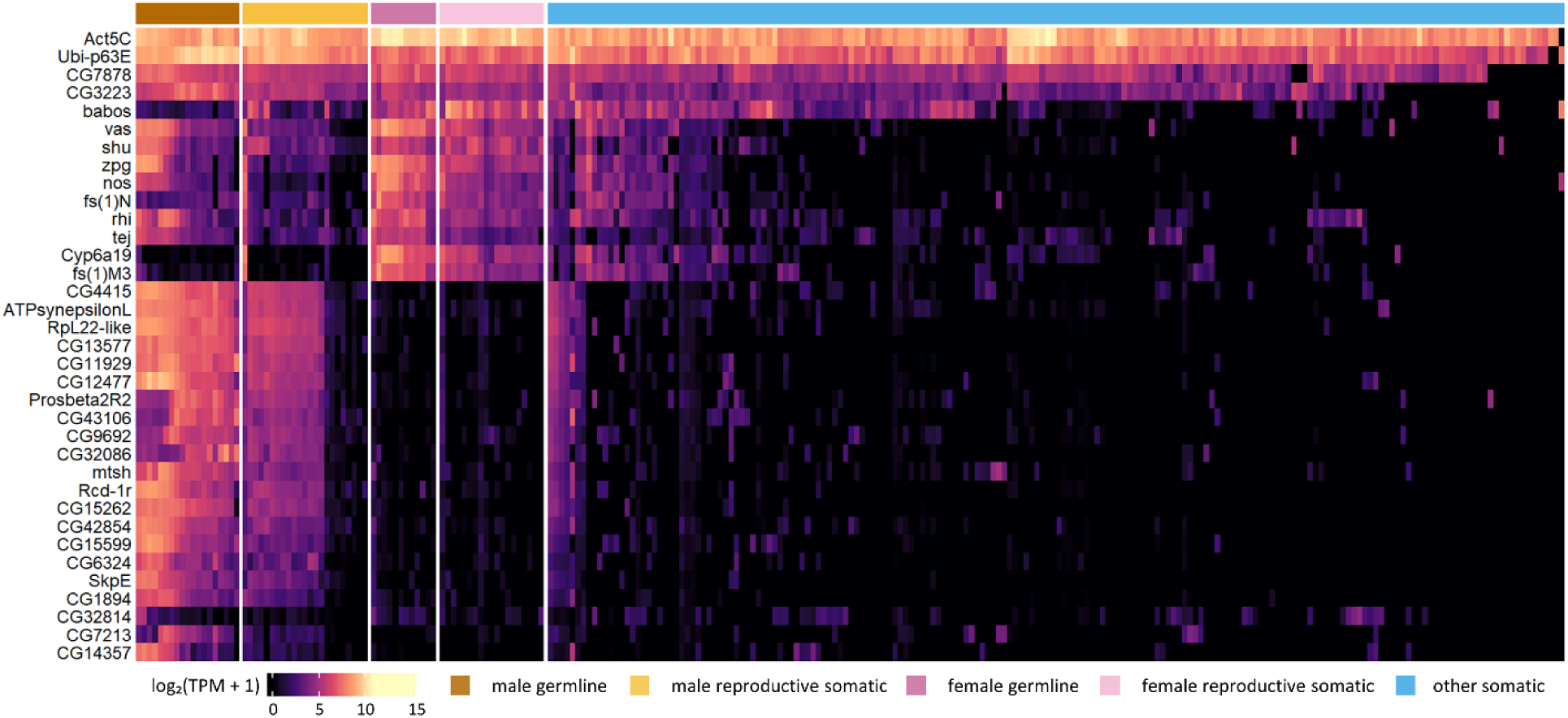
Single cell expression profiles of promoter candidate genes in *Drosophila melanogaster*. The heatmap displays the expression profiles of 35 gene candidates across 259 single cell subtypes from the Fly Cell Atlas. Raw Transcripts Per Million (TPM) were transformed to log_2_(TPM + 1) and color-mapped. To determine gene and cell type order, we computed Euclidean distances on the transformed expression matrix and performed complete-linkage hierarchical clustering.

### Different promoters exhibit substantially varied drive performance

We systematically evaluated the drive performance of various Cas9 promoters within a split-drive system targeting the *yellow* gene on the X chromosome. In this system, the gRNA directs Cas9 to cleave *yellow*, enabling HDR to convert the wild-type allele into a drive allele (Fig.2A). We assessed the drive performance of different Cas9 promoters by performing crosses and analyzing the progeny based on the fluorescence markers and body color phenotypes (Fig.2B, Fig.S2C).

**Fig. 2:**
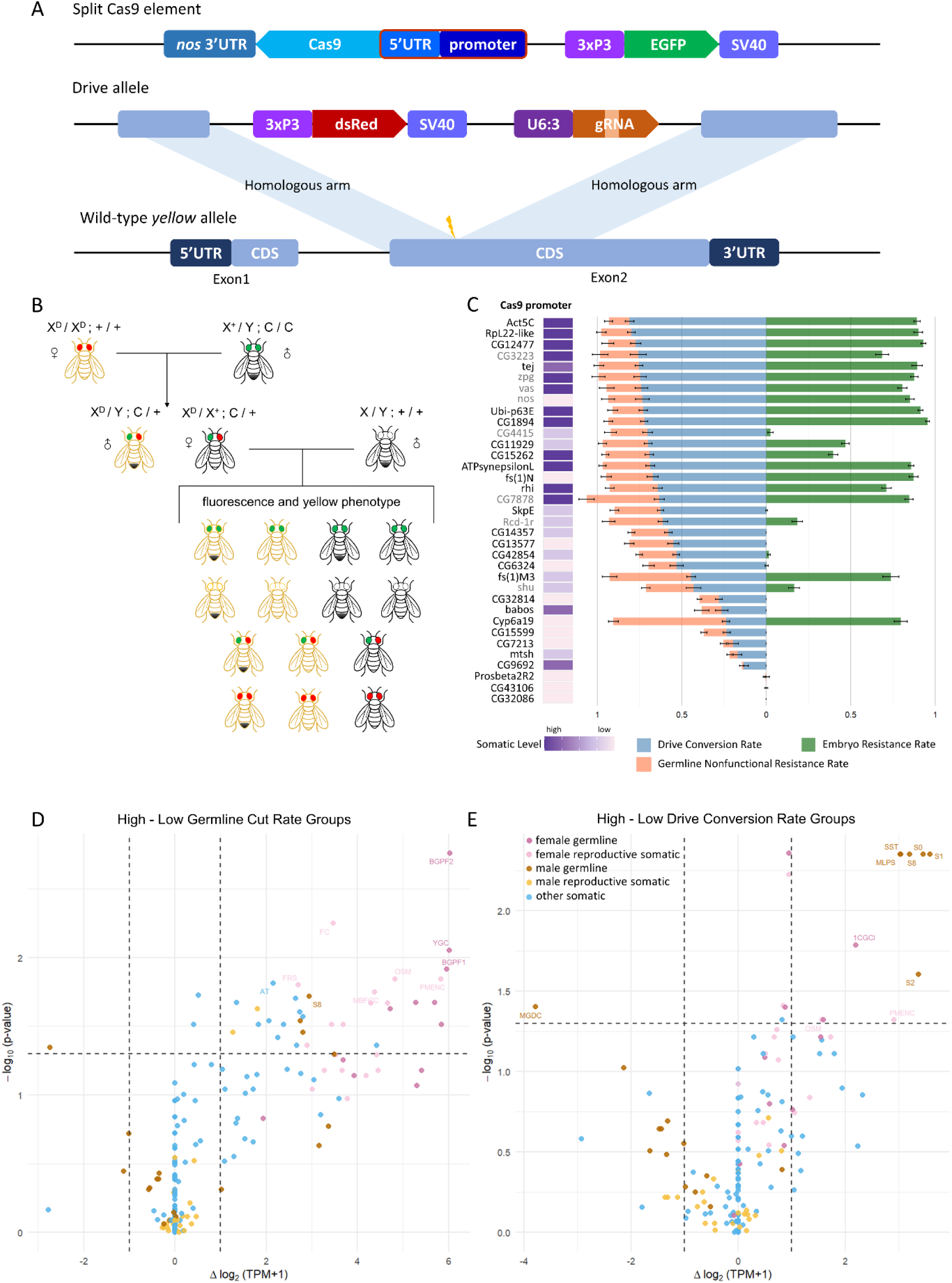
Performance of Cas9 promoters with the split drive targeting *yellow*. **A.** The *yellow* system includes two components: a split Cas9 element and the drive element. The drive element contains a DsRed fluorescent marker under the control of the 3×P3 promoter and a gRNA driven by the U6:3 promoter, targeting *yellow* on the X chromosome. The split Cas9 element is located on chromosome 2R and expresses Cas9 under the control of different promoters and the *nanos* 3′ UTR, along with an EGFP marker also driven by the 3×P3 promoter. The gRNA in the drive element directs Cas9 to cleave exon 2 of *yellow*. **B.** The cross scheme of the split drive targeting *yellow* on the X chromosome. Females with the drive element strain are first crossed to males with the Cas9 element. Females carrying both fluorescent markers are selected from the progeny, and their body color is used as an indicator of somatic Cas9 activity. Drive and Cas9 females were then crossed with *w^1118^* males, and their offspring was phenotyped to assess drive performance. **C.** Drive performance of 35 Cas9 constructs with promoters. The sum of the drive conversion rate and the germline nonfunctional resistance rate is close to the total germline cut rate. Data from grey labeled Cas9 promoters are from a previous study^36^. For visualization purposes, drive conversion rates calculated as below zero are displayed as zero (these are statistically indistinguishable from zero). Error bars indicate standard error of the mean. Source data are provided in Data Sets S1-S2. **D.** Differential expression of promoter-associated genes between high and low cut rate groups across various cell types. Each point represents a single cell type from the Fly Cell Atlas dataset, with the horizontal axis indicating the difference in log_2_(TPM+1) between high and low cut rate groups, and the vertical axis showing -log_10_(*p*-value) derived from Wilcoxon rank-sum tests. Top cell types with significant differences (*p* < 0.05 and |Δlog_2_(TPM+1)| > 1) are labeled with their abbreviated names (Table S2). Source data are provided in Data Set S5. **E.** Differential expression of promoter-associated genes between high and low drive conversion groups across various cell types. Plot structure and interpretation as in part D. Source data are provided in Data Set S6.

The constitutive promoters *Act5C* and *Ubi-p63E* display exceptionally high somatic expression, consistent with their characterization as ubiquitously active promoters (Fig.2C, Fig.S3A). However, both promoters also exhibited unexpectedly high drive conversion of 81% and 73%, respectively. The embryo resistance formation was high at 89% and 91%, respectively. Seventeen other germline promoters also showed high drive conversion rate (over 60%). Among them, the *CG4415* and *SkpE* promoters also exhibited low embryo resistance rate, and *Rcd-1r*, *CG11929*, and *CG15262* exhibited moderate levels of embryo resistance formation. Six promoters exhibited moderate drive conversion rate (ranging from 30% to 60%), with *CG6324*, *CG42854*, *CG13577*, and *CG14357* also having extremely low embryo resistance. Seven promoters showed relatively low drive conversion rate (10% to 30%), among which only *Cyp6a19* displayed high total germline cut rate (it also had high embryo resistance). The remaining six promoters showed moderate to low germline cut rates and did not generate embryo resistance alleles. Additionally, three promoters did not exhibit any detectable cut activity (Fig.2C, Fig.S3A).

To understand these results in the context of transcriptome profiles, we performed a systematic correlation analysis between drive performance and the predicted expression patterns of each promoter, as inferred from the Fly Cell Atlas. Among promoters with high total germline cut rate (>90%), the five cell types showing the most elevated expression were all female reproductive system cells, four of which were germline (Fig.2D, Fig.S4). Several other female germline and non-germline cells showed moderate elevation, and a few cells of other types showed small but statistically significant changes in expression (usually increased expression). Because our total cut rate in this X-linked system was for females, these results are consistent with expectations. We also performed a similar analysis focusing on the drive conversion rate. Compared to promoter genes with low drive conversion (less than 15%), those associated with high drive conversion (more than 40%) showed significantly higher expression across 15 reproductive system cells in *Drosophila*, comprising eight female and seven male cell types, with male germline cells showing the highest levels of effect size and statistical significance (Fig.2E, Fig.S5). Because *yellow* is located on the X chromosome and drive conversion rates can only be measured in female individuals, we hypothesize that the statistical significance observed in male germline cells may be due to co-regulation of certain genes in both male and female germline cells, which together with limited signal size and uncertain female germline cell classification may have resulted in male germline cells yielding the strongest signal in this analysis.

### High drive conversion in different sexes is associated with elevated expression in the corresponding germline cells

To compare drive performance between males and females and potentially examine co-regulatory effects of promoters in male and female germline cells, all promoters were tested in another drive system that allows assessment of drive conversion rate in both sexes. This system is also a split gRNA element, but targeting *cinnabar* located on chromosome 2 (Fig.3A). Drive conversion associated with each Cas9 promoter was assessed through crosses based on fluorescence phenotypes (Fig.3B-C), but embryo resistance could not be assessed for these because all tested individuals had a *w^1118^* background.

**Fig. 3:**
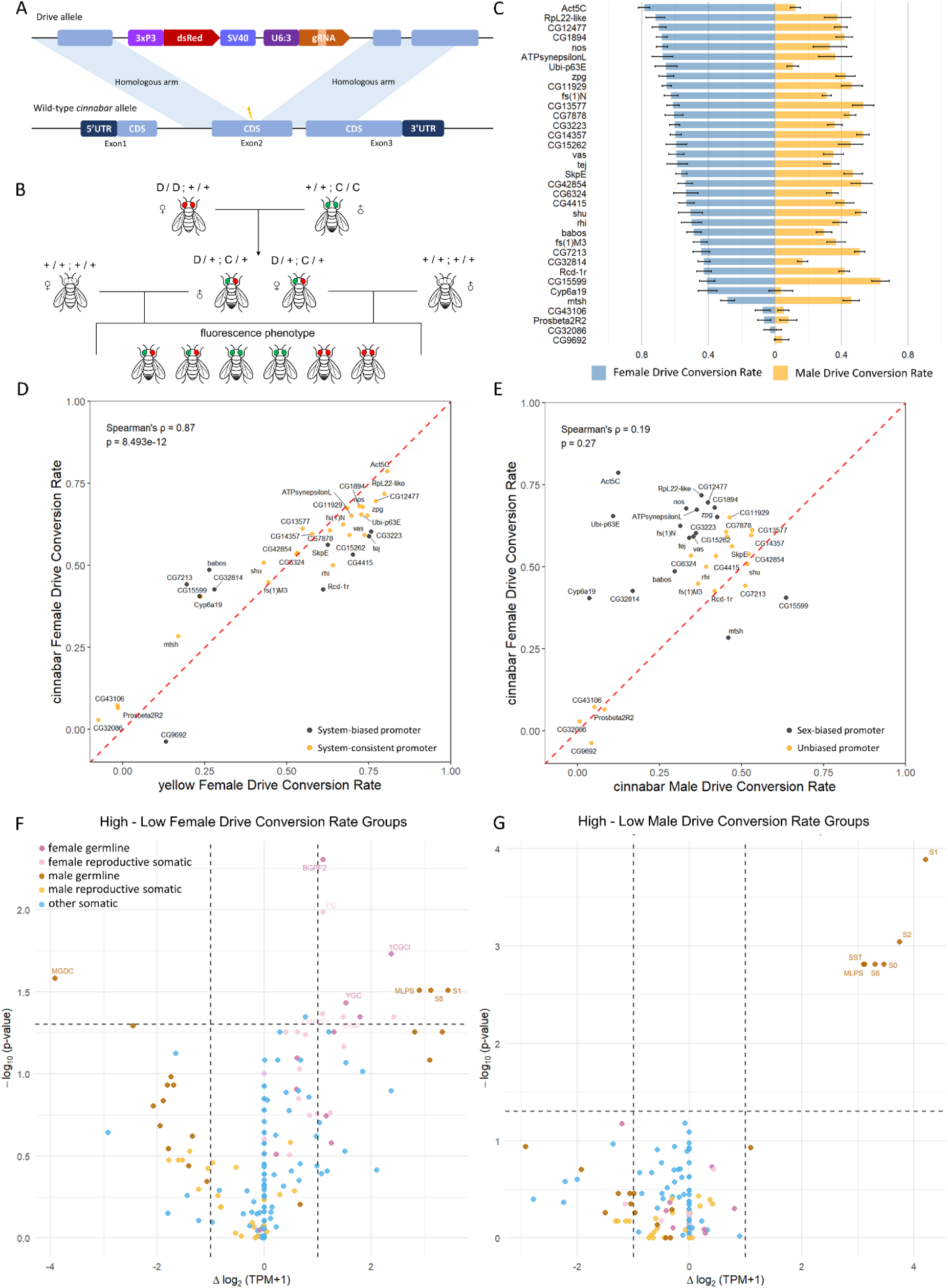
Performance of Cas9 promoters with the split drive targeting *cinnabar*. **A.** The *cinnabar* system includes two components: a split Cas9 element and the drive element. The gRNA in the drive element directs Cas9 to cleave exon 2 of *cinnabar*. **B.** The cross scheme of the split drive targeting *cinnabar*. Females from the drive element strain are first crossed with males from the Cas9 strain. Females or males with both drive and Cas9 are selected from the progeny and crossed to *w^1118^*individuals. Drive performance was assessed by analyzing the phenotypes of the resulting offspring. **C.** Drive performance of 35 Cas9 constructs regulated by different gene promoters in the *cinnabar* system. For visualization purposes, calculated drive conversion rates below zero are displayed as zero (these are statistically indistinguishable from zero). Error bars indicate standard error of the mean. Source data are provided in Data Set S3. **D.** Comparison of female drive conversion rates between the *cinnabar* and *yellow* drive systems. Each point represents a promoter construct. Orange points indicate promoters showing no significant difference in drive conversion rate between the two systems (*p* ≥ 0.05, based on t-test or Wilcoxon test). Spearman correlation analysis revealed a strong positive association between the two systems (ρ = 0.87, *p* = 8.49×10^-12^). **E.** Comparison of female and male drive conversion rates in the *cinnabar* system. Each point represents a promoter construct. Spearman correlation analysis revealed a moderate positive association between male and female drive conversion (ρ = 0.19, *p* = 0.27). **F.** Differential expression of promoter-associated genes between high and low female drive conversion groups across various cell types. Each point represents a single cell type from the Fly Cell Atlas dataset, with the horizontal axis indicating the difference in log_2_(TPM+1) between high and low female drive conversion groups, and the vertical axis showing the -log_10_(*p*-value) derived from Wilcoxon rank-sum tests. Most cell types with significant differences ( *p* < 0.05 and | Δlog_2_(TPM+1)| > 1) are labeled with their abbreviated names (Table S2). Source data are provided in Data Set S7. **G.** Differential expression of promoter-associated genes between high and low male drive conversion groups across various cell types. Plot structure and interpretation as in part F. Source data are provided in Data Set S8.

For female drive conversion, the results obtained from the *cinnabar* system were highly consistent with those from the *yellow* system (Fig.3D), indicating that the influence of promoters on drive performance is likely generalizable across different gene drive systems, at least in most cases. Moreover, promoters that exhibited high drive conversion in males often showed similarly high conversion in females (Fig.3E), supporting the existence of a co-regulatory effect between the sexes. We also investigated other promoters with higher female-bias in drive conversion. Compared with promoter-associated genes having unbiased drive conversion, genes with female-biased drive conversion exhibited higher expression in female reproductive system cells (Fig.S6). This could potentially be because male and female germline cells require different expression levels and/or timing to achieve the same drive conversion rate.

Also similar to the *yellow* system, the promoters with high drive conversion rates in females typically displayed significantly elevated expression in female and male germline cells (Fig.3F), as exemplified by *nanos*, *vasa*, and *fs(1)N*, which both showed high expression in ovary cells corresponding with high female conversion rate (Fig.S7, Fig.S8A-C). In males, the pattern was simpler, with only male germline cells displaying significant differences, predominantly early male pre-meiotic germline cells (Fig.3G). *CG15599*, *CG12477*, and *CG14357* exemplify this, showing high expression in testis cells and high male conversion rate (Fig.S8D-F. Fig.S9). These findings suggest that high drive conversion in each sex remains closely associated with elevated expression in the corresponding germline cells.

However, we noted exceptions such as *Act5C* and *Ubi-p63E*, which showed high expression in all cell types of both sexes but did not have high drive conversion in males (Fig.3C), despite high drive conversion in females (Fig.2C, Fig.3C). To investigate this discrepancy, we performed target site sequencing on progeny derived from crosses between drive males and the wild-type females, focusing on offspring that did not inherit the drive. The results showed that all target sites in these progeny were successfully cleaved by Cas9, indicating that the low conversion rates in males were not due to inefficient cleavage, but were more likely a consequence of repair bias away from HDR and toward end-joining (Fig.S10). This observation suggests that high germline expression alone is not always sufficient for efficient drive conversion, highlighting the importance of considering the HDR efficiency and timing within the germline cells of each sex. It is possible that these ubiquitously active promoters may have induced cleavage in germline stem cells earlier than our germline promoters, and cleavage at this time in males (but not females) could be less likely to be repaired by HDR.

### Undesirable drive performance is associated with excessive expression of Cas9

Undesirable drive performance is characterized by limited drive conversion, accumulation of resistance alleles, and fitness cost to carriers. Two key contributors are embryo resistance allele formation and somatic Cas9 expression. Resistance alleles formed in the zygote or early embryo through end-joining can block subsequent cutting and conversion, creating a major barrier to drive spread, especially when the drive itself imposes fitness costs. For suppression drive, this can also lead to premature removal of drive alleles in sterile drive/nonfunctional resistance allele heterozygotes. In addition, somatic Cas9 activity in suppression drives (and some modification drive variants) imposes fitness costs that reduce reproductive competitiveness, ultimately limiting drive power.

To investigate the relationship between resistance allele formation and Cas9 expression patterns, we compared the transcriptomic profiles of genes corresponding to promoters in the *yellow* drive system that exhibited drive conversion rate higher than 40% (representing potentially viable drive candidates), along with either high embryo resistance rates (more than 65%) or low rates (less than 5%). Based on bulk RNA-seq data, differential expression analysis revealed that the formation of embryo resistance alleles is primarily associated with expression in early embryo stages and female adults, as expected (Fig.S11). Additionally, analysis of single-cell transcriptomic data revealed that the formation of embryo resistance alleles is strongly associated with excessive Cas9 expression in female germline cells (Fig.4A, Fig.S12). This is congruent with the notion that embryo resistance allele formation is the result of Cas9 deposition in gametes, which is then transmitted to the zygote. Male-specific cells had no substantial correlation with embryo resistance except for unknown testis cells, which may be related in their expression pattern to female cells, or perhaps include some precursor cell types shared by both sexes.

**Fig. 4:**
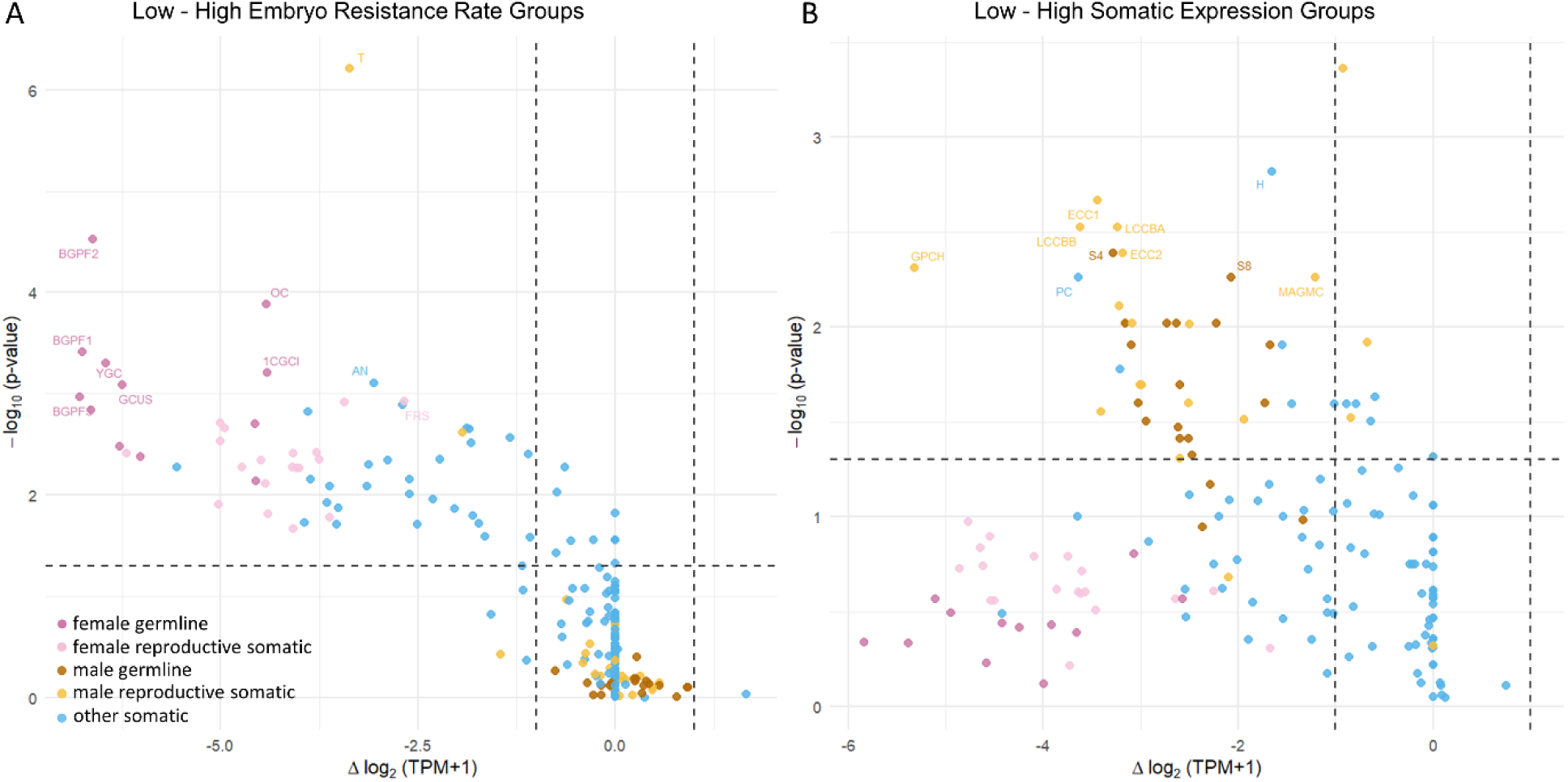
Differential cell type expression analysis of promoter associated genes for high and low embryo resistance and somatic activity. **A.** Differential expression of promoter-associated genes between low and high embryo resistance rate groups across various cell types. Each point represents a single cell type from the Fly Cell Atlas dataset, with the horizontal axis indicating the difference in log_2_(TPM+1) between high and low embryo resistance rate groups, and the vertical axis showing the -log_10_(*p*-value) derived from Wilcoxon rank-sum tests. Select cell types with significant differences (*p* < 0.05 and |Δlog_2_(TPM+1)| > 1) are labeled with their abbreviated names (Table S2). Source data are provided in Data Set S10. **B.** Differential expression of promoter-associated genes between low and high somatic expression groups across various cell types. Plot structure and interpretation as in part A. Source data are provided in Data Set S11.

A similar analysis of promoters associated with high and low somatic Cas9 expression (Fig.S2) showed that somatic expression from our promoters is not necessarily strongly linked to excessive Cas9 expression in somatic cells, though a few somatic cells types had significant correlation (Fig.4B, Fig.S13). Unexpectedly, the majority of significant cell types were associated with the male reproductive system, where high expression was negatively correlated with somatic expression. This may be in part due to our selection of promoters that were mostly germline restricted. However, the lack of statistical significance for female cells (despite slightly higher effect size) may well still hold scientific meaning. It may be that when promoters are used with non-native genes such as Cas9 at non-native genomic sites, then male expression patterns are more easily lost, with additional somatic “leakiness” where none existed before. It is also possible that this phenomenon is due to our specific chosen Cas9 genomic locus.

To further assess somatic expression, we used a *doublesex* suppression system^54^ to examine the Cas9 promoter constructs that exhibited low somatic expression in the *yellow* system (Fig.S2). None of the females with both drive and Cas9 displayed a strong intersex phenotype, which was previously indicative of somatic expression^50^. We also found that these females did not exhibit sterility at greater rates than wild-type (though we cannot rule out small to moderate fertility reductions), even with lines that had small amounts of visible somatic activity in the *yellow* system. These results show that several of our Cas9 constructs maintained consistently relatively low somatic expression between systems, though small differences in expression timing and location may somewhat change fitness effects between different targets.

### Ideal drive performance requires a narrow range of Cas9 expression in female germline cells

Ideal drive performance is characterized in part by a combination of high drive conversion rate, low embryo resistance allele formation, and low leaky somatic expression. Investigating the link between ideal drive performance and single-cell transcriptomic expression patterns in the female germline, we found that female drive conversion is associated with high Cas9 expression in certain female germline cell types, but low embryo resistance allele formation is linked to low Cas9 expression in those same cell types (Fig.5A-B). This suggests that achieving ideal drive performance may require Cas9 expression to be maintained within a specific expression window in particular cell types, which could include a temporal window and/or a quantitative window. To further assess this, we examined the expression profiles of genes corresponding to three kinds of promoters with distinct drive performance, focusing on their expression in female germline cells. Results indicate that optimal drive performance is achieved only when promoter activity is tightly constrained within a narrow expression range in these cell types (Fig.5C). While this does not rule out a wider expression quantity window if expression timing was further optimized, it does at least indicate that achieving good performance is possible, even if difficult.

**Fig. 5:**
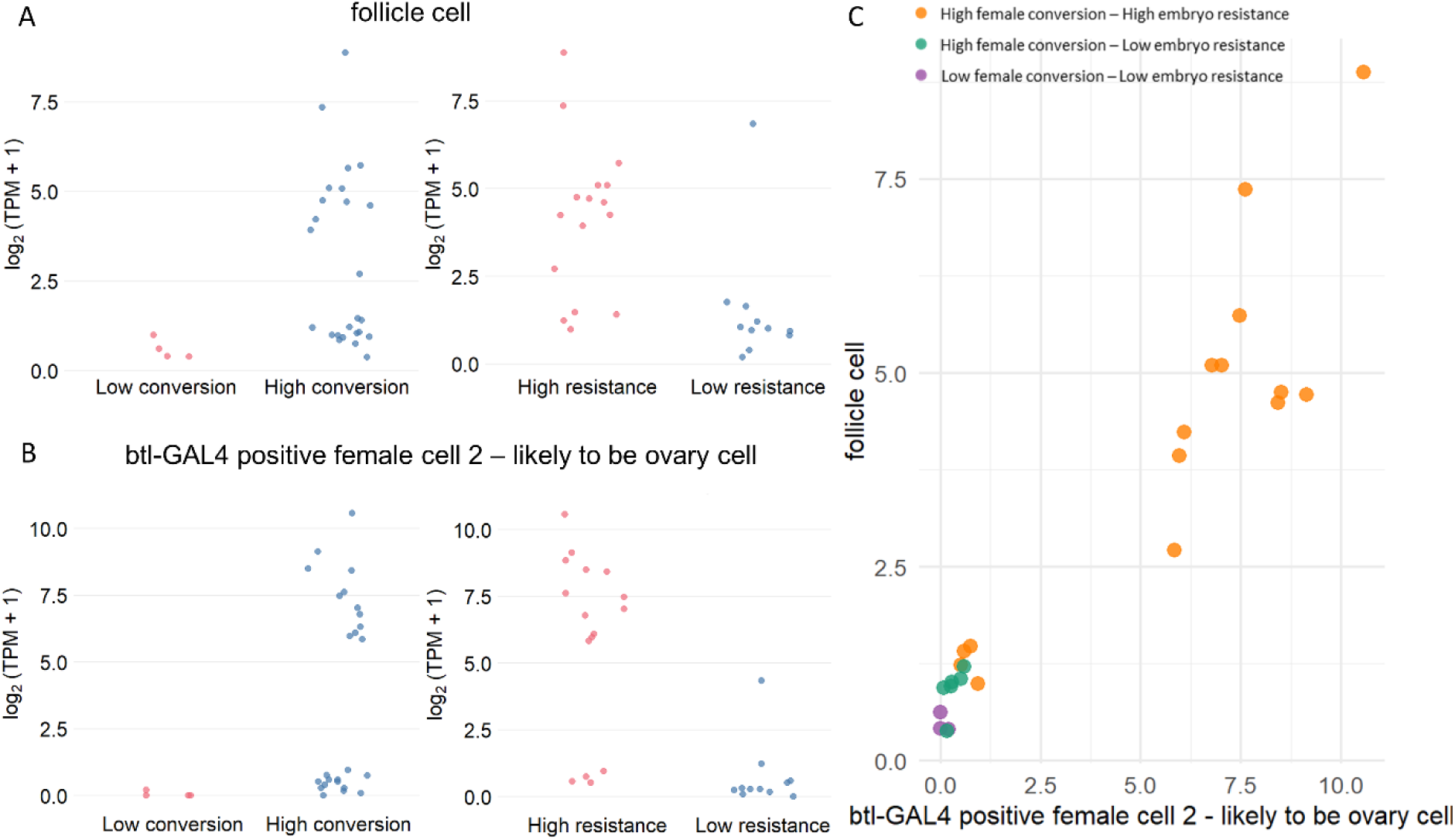
Expression profiles of promoter-associated genes in female germline cells for optimal drive performance. **A.** Expression differences of promoter-associated genes in follicle cells. The left panel compares high conversion promoters with low conversion promoters. The right compares high resistance with low resistance. The vertical axis shows log₂(TPM + 1) expression of the genes in follicle cells. Each dot represents the expression value of one promoter-associated gene. Statistical differences were evaluated using the Wilcoxon rank-sum test ( *p* < 0.05 considered significant). **B.** Expression differences of promoter-associated genes in btl-GAL4 positive female cell 2 - likely to be ovary cell. Plot structure and interpretation as in part A. **C.** The range of Cas9 expression in female germline cells. The plot shows log₂-transformed expression levels of selected promoter-associated genes in two female reproductive cell types: btl-GAL4 positive female cell 2 – likely to be ovary cell and follicle cell. Promoters are grouped into three categories based on their drive performance in the *yellow* system. Each dot represents one promoter.

Based on the results in the *yellow* and *cinnabar* drive systems, we selected six Cas9 promoter constructs with superior drive performance and evaluated their drive inheritance in a drive system targeting *RpL35A*. This drive was previous found to be effective at removing resistance alleles, yielding high drive inheritance and low to moderate embryo resistance allele formation, but it also had lower germline cut rates in previous tests^8^, perhaps only just enough to support high drive conversion. All six of our new Cas9 constructs exhibited high drive inheritance in males, whereas female drive inheritance was comparatively low (Fig.S14). This is consistent with our transcriptomic analysis, which shows that a narrow activity window is necessary for good drive inheritance but low embryo resistance allele formation in females. This lower activity drive element was close to this ideal window with the high expression *nanos* promoter, which had too high of an embryo resistance rate in the *yellow* system. However, for our promoters that had ideal activity in the *yellow* system, their activity was too low in the *RpL35A* system. It is interesting that they still maintained high activity in males, a contrast to our investigation in *cinnabar* where there were no candidates that had low activity in females but high activity in males.

### *In situ* expression of Cas9 at the *rhino* locus prevents leaky somatic expression

Gene drives usually require use of native gene promoters at non-native sites. In our investigation, we kept the genomic locus constant to facilitate comparison between promoters. Some differences were seen between drive performance and expected native gene expression patterns (Fig.1-3). This could potentially be due to different genomic locus or due to missing regulatory elements in synthetic promoters. We also used the *nanos* 3ʹ UTR due to superior performance in previous tests^36^, but this could have influenced our new promoters in different ways. To investigate how regulatory sequences of different lengths and genomic contexts shape Cas9 expression and drive performance, we designed four Cas9 expression constructs (RO1, RO2, RO3, and RON - only RO3 was used in the analysis in previous sections) derived from the regulatory regions of the *rhino* (*rhi*) gene. RO1, RO2, and RO3 were inserted into our usual neutral site on chromosome 2R and carried 441 bp, 1373 bp, and 3422 bp of the upstream *rhino* regulatory sequence, respectively (Fig.6A). RO2 excluded the coding sequence of a nearby gene and was thus composed of two separate promoter elements. Transcription termination was mediated by the *nanos* downstream regulatory sequence in RO1 and RO2, and by *rhino* in RO3. The RON construct was generated by inserting Cas9 directly into the endogenous *rhino* locus (together with a copy of the *rhino* 3ʹ UTR), thereby enabling it’s *in situ* expression (Fig.6B). This construct disrupted the native *rhino* gene, causing recessive female sterility^48,49^, so only male carriers instead of male homozygotes were used to generate flies for testing.

**Fig. 6:**
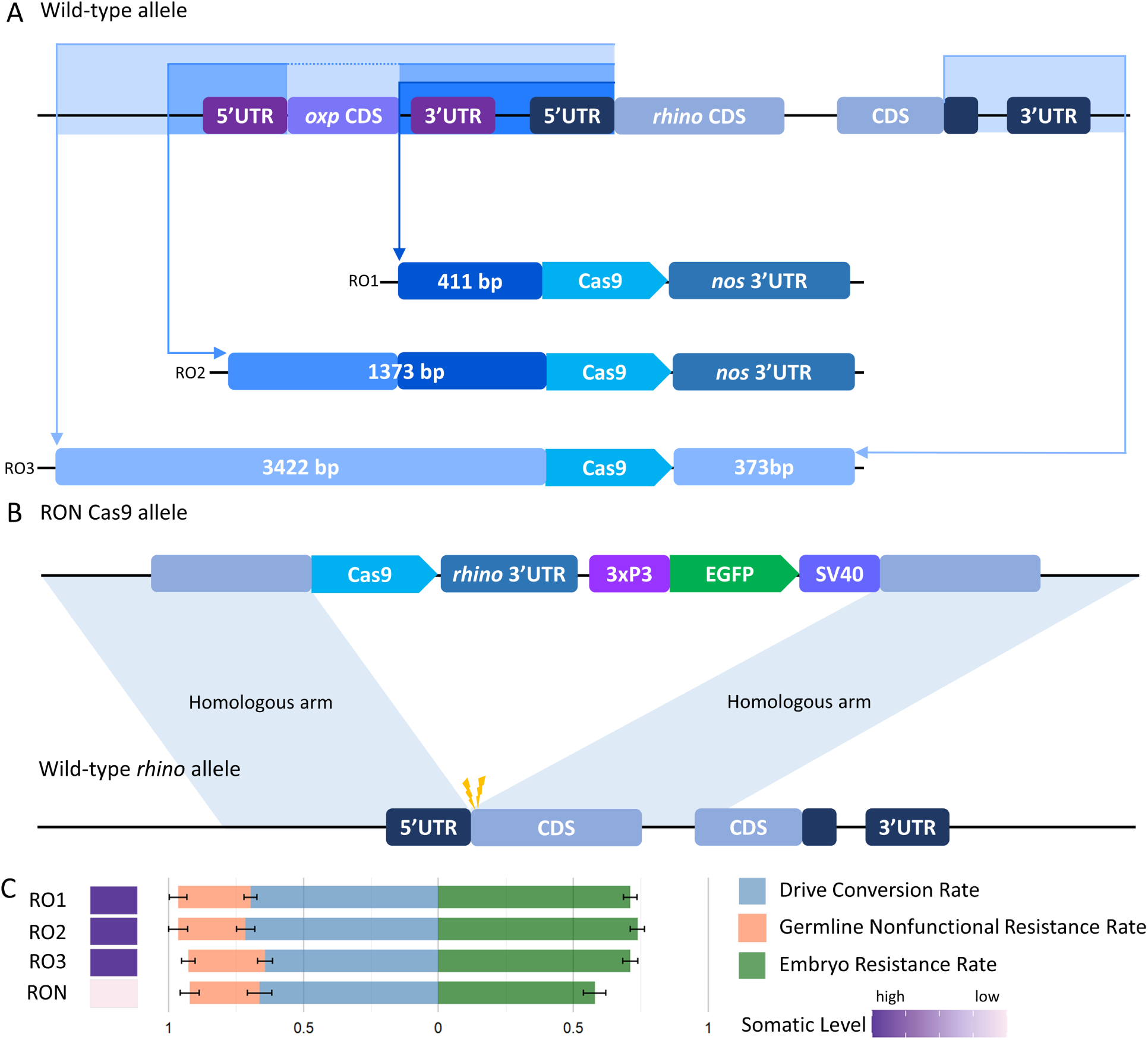
Drive performance of four Cas9 constructs with different *rhino* regulatory regions. **A.** Three ectopic Cas9 constructs based on *rhino*. These were integrated into a neutral site on chromosome 2R and carried EGFP as a fluorescent marker. They contained different lengths of upstream *rhino* regulatory sequence. The downstream regulatory sequence was derived from *nanos* for RO1 and RO2 and from *rhino* for RO3. **B.** An *in situ* Cas9 construct based on the *rhino*. This construct carried EGFP and Cas9, but it was inserted directly into the *rhino* locus, with Cas9 replacing the *rhino* coding sequence immediately after the 5′ UTR. **C.** Drive performance of the Cas9 constructs in the *yellow* system. Error bars indicate standard error of the mean. Source data are provided in Data Sets S1-S2.

Drive performance evaluated through the *yellow* system revealed that the three ectopic Cas9 constructs showed no significant differences in any aspect of drive performance (Fig.6C). This suggests that promoter element length itself was not a determining factor, as long as the critical regulatory regions are present. In addition, use of the native 3′ UTR did not noticeably alter drive performance. However, the *in situ* expression construct showed a slight decrease in embryo resistance rate and exhibited markedly reduced Cas9 activity in somatic cells compared to the three ectopic expression constructs. This finding indicates that when Cas9 is expressed *in situ* at the *rhino* locus, its transcription and translation are subjected to tighter endogenous regulation, thereby reducing leaky expression, especially in somatic cells.

## Discussion

In this study, we evaluated 35 Cas9 lines driven by different promoters across multiple types of gene drive systems. By comparing their performance within these systems, we not only identified promoters with good drive performance, but also, through integration with single-cell transcriptomic data, found potential associations between drive performance and promoter properties. Previous studies have suggested that restricting Cas9 expression to germline cells using germline-specific promoters can substantially improve drive performance. Our findings support this notion. Furthermore, our results showed that drive conversion in each sex correlates with the expression level of the promoter-associated gene in the corresponding germline cells. In addition, embryo resistance allele formation is associated with excessive Cas9 expression in female germline cells, while somatic activity is associated with male germline cell expression of native genes. Taken together, these findings reveal that optimal drive performance is strongly associated with a narrow range of Cas9 expression in female germline cells. Though exact quantitative levels may differ across species, this pattern could potentially allow for easier selection of Cas9 promoters in non-model organisms while yielding insights into drive performance complexities.

We evaluated Cas9 promoters across multiple gene drive systems. Initially, we performed a preliminary screen using the *yellow* system^7^. To enable assessment of male drive performance, we then employed the *cinnabar* system. Finally, the promoters with superior drive performance were assessed in well-studied and more advanced drive systems targeting *doublesex*^50^ or *RpL35A*^8^, representing suppression and modification drives, respectively. This strategy demonstrated that Cas9 promoter performance can be at least somewhat generalized across distinct drive systems, even if gRNA activity levels and genomic location also play an important role in drive performance.

For the selection of promoter regulatory regions, we identified regions based on transcription factor binding densities from the ReMap database^53^. However, this approach cannot guarantee complete coverage of all regulatory elements, in particular distal regulatory elements (enhancers)^55,56^, features mediated by higher-order chromatin structures^57^, and noncanonical regulatory modes (such as epigenetic mechanisms)^58^. Despite these limitations, integrating our evaluation of drive performance across multiple drive systems with single-cell transcriptomic analyses indicates that for most cases, the selected regions mechanistically account for differences in drive conversion, resistance allele formation, and leaky somatic expression, and align with the expression levels and temporal patterns of the promoter associated genes in certain cell types. It remains possible that our low-activity promoters in particular may be missing important enhancers. Taken together, though, these observations suggest that the promoter regions included in this study at least mostly capture the critical regulatory elements of their associated genes.

For the selection of 3′ UTR, we uniformly used the *nanos* 3′ UTR rather than the endogenous 3′ UTR corresponding to each promoter. This decision was based on previous findings showing that the *nanos* 3′ UTR not only lacks observable negative effects but may, in some cases, enhance drive performance^36^. Using a consistent 3′ UTR while varying only the promoter allowed us to isolate and evaluate the effects of different promoters. However, gene expression regulation is not solely dependent on the promoter and 5′ UTR^59^. Several studies have shown that 3′ UTRs possess regulatory capabilities comparable to those of promoters^60–62^. Moreover, some endogenous genes have been proven unable to be expressed through promoter activation alone, but instead require simultaneous replacement of the 3’ UTR^63^. These findings suggest that using both the corresponding promoter and 3’ UTR sequences should better approximate expression patterns of endogenous genes. It is possible that some of our promoters did not function or functioned differently due to use of the *nanos* 3′ UTR. Yet, this study and previous gene drives studies did not show substantially different results when the 3′ UTR was varied, so it remains unclear under what circumstances the exact form of the 3′ UTR becomes important.

In this study, most Cas9 constructs were inserted into a neutral site on chromosome 2R, downstream of both adjacent genes. Notably, our results showed that our *rhino* promoter *in situ* construct, while maintaining drive performance comparable to the ectopic constructs, markedly reduced somatic Cas9 activity. This finding further underlines the importance of endogenous spatiotemporal regulation. The *in situ* construct, by relying on the gene’s own endogenous regulatory framework, achieved more precise control of Cas9 across developmental stages and tissue specificity, thereby effectively avoiding leaky expression in non-target cells. This observation suggests that *in situ* expression could not only serve as a strategy to reduce the fitness costs associated with somatic expression but also provide a potential means to further enhance the drive performance. Self-limiting split drive strategies and particularly the integral drive strategy^64,65^ are well-placed to take advantage of this finding, though there may be increased complexity in overcoming challenges with resistance alleles and native gene expression as a tradeoff for this. Most gene drives, though, require the Cas9 gene to be at a non-native locus or cannot take advantage of this strategy for other reasons (for example, suppression drive targets using native regulatory elements to express Cas9 may be cleaved and lose essential expression in heterozygotes unless the delay in CRISPR cleavage is sufficiently long for enough native gene product to first be expressed). It remains unclear whether the native locus in our test had specific regulatory elements or genomic structure that may interact with Cas9 expression or if our chosen site for most of our testing itself makes genes more prone to somatic activity. If it is the latter, then prospects for achieving optimal drive activity based on single-cell transcription data are improved because there will usually be multiple possible genomic sites and target genes for drive designs.

During the selection of candidate genes, we attempted to choose promoters with sex-specific expression patterns based on transcriptomic profiles (Fig.1, Fig.S1). For example, *fs(1)M3* and *babos* were found to be highly expressed in female germline cells, whereas *CG14357* and *CG7213* showed high expression in male germline cells. However, drive performance assays revealed that these promoters exhibited relatively high conversion in both male and female germline cells (Fig.2C). Transcriptome analysis further revealed that these genes were not entirely silenced in the germline cells of the opposite sex, but it remains surprising that their expression was sufficient for strong drive activity, in line with Fig.5C showing that promoter candidates with superior drive performance exhibit only low expression in two significant female reproductive cell types.

It is noteworthy that although *Act5C* and *Ubi-p63E* are widely regarded as strong constitutive promoters with similar expression in most cell types, they exhibited marked sex-specific differences in the *cinnabar* drive system. We thus hypothesize that the timing of HDR-biased repair or perhaps Cas9 cleavage with the same promoters differs between male and female germline cells, affecting the outcome of drive conversion. In females, Cas9-mediated cleavage occurring during early germline development is perhaps more likely to engage the HDR pathway, thereby enabling successful drive conversion. In contrast, similar cleavage events in males may be prone to repair through end-joining, resulting in the formation of resistance alleles. This observation implies that the time window for HDR in male germline cells may be later than in females (and note also that expression in early male germline cells appeared to be optimal). Further experimental evaluation may resolve this. Perhaps related to this, the identity of male germline cells in the single-cell transcriptome data has higher confidence that female germline cell types, likely due to much higher quantity and other associated developmental factors. Gaining greater resolution in female cell types may thus allow for improvement in female transcriptomic data, potentially yielding information that could clarify this situation and allow for greater effectiveness when applied for gene drive optimization.

We unexpectedly observed that leaky somatic expression is correlated with high expression of a class of promoter-associated genes in male reproductive system cells. This phenomenon could potentially be attributed to biases introduced during promoter selection. First, in single-cell transcriptomic (scRNA-seq) data, male reproductive system lineages have more granular and consistent developmental stage annotations, whereas the annotation of female lineages is less complete than the former. Consequently, we were more likely to identify and select male promoters with desired expression patterns, resulting in an inflated proportion of male-specific and possible introduction of selection bias. In addition, this limitation influenced our significance analyses. We could pinpoint the optimal stage for male drive conversion with good precision (early pre-meiotic germline cells), whereas for females, we could only could only implicate some early germline cell types and could not confidently assign specific developmental stages to the remaining female germline cells (Fig 3F, G). Secondly, our selection strategy prioritized promoters exhibiting stage-specificity and high expression within different germline cells. Although this strategy provides a more precise basis for evaluating and comparing drive performance across developmental stages, it also decreases the overall somatic expression in our candidates compared to most other genes. However, the bias described above mainly influences the analysis of somatic expression and is unlikely to exert an impact on the evaluation of other drive performance analyses. The principal reason for this is that these two analyses operate at different levels and are statistically stratified: drive performance such as drive conversion rate and resistance formation rate are evaluated by the relative comparisons in reproductive system lineages, whereas somatic expression is determined primarily by the somatic cells from other tissues. Expansion of promoter candidates to include specificity from multiple somatic organs could further refine our analysis of somatic expression and thereby establish a more balanced reference and improving the accuracy of the analysis.

We acknowledge that the actual expression pattern of Cas9 may not fully replicate *in situ* expression of the corresponding genes, but our experimental results nonetheless show that in most cases, they are closely correlated. This indicates that our approach should be broadly applicable despite individual exceptions. Based on the results of this study, we propose a workflow for screening Cas9 promoters across different species to enable efficient Cas9 expression in gene drive systems. First, the required expression activity and spatiotemporal window for Cas9 should be clearly defined according to the mechanism of the designed drive system. For example, in suppression drives, it is necessary to achieve high Cas9 expression in germline cells to enable efficient cleavage at target sites while avoiding somatic expression as much as possible to reduce fitness costs caused by off-target cleavage. Embryo resistance should also be avoided, though this is not as high priority as avoiding somatic activity. In modification rescue drives, embryo resistance is still harmful, but it is more important to maximum drive conversion. Somatic activity is substantially less harmful than for suppression drives. A notable exception to this is drives targeting haplolethal genes, where embryo resistance and somatic expression are harmful to the drive (though with their priorities are probably reversed compared to suppression drives), but potentially still worth the improved rate of nonfunctional resistance allele elimination.

After obtaining high-quality scRNA-seq data or bulk RNA-seq data for the species of interest (which has already been at least partially conducted for some major vectors^66,67^), a screening strategy should be formulated based on the drive requirements. For species with scRNA-seq data, the requirements should first be mapped to the corresponding cell clusters. High drive conversion typically requires medium to high levels of expression in germline cells at early developmental stages, while the low embryo resistance rate favors low or absent expression in germline cells at later developmental stages or in embryos. There is a narrow window where both of these can be achieved. However, it should be noted that this window may be quantitatively different for different species and drives. For example, variance in specific gRNA activity may have allowed a low-expression system to produce good drive conversion in our *yellow* system, but higher activity would be needed for other less active gRNA targets such as our *RpL35A* system to achieve the same total cut rate. To avoid somatic expression, low or absent expression in somatic cells is required, and such absence of somatic activity may be correlated with low expression in male reproductive system cells. The desired expression specificity of each gene in germline cells should then be assessed, prioritizing genes with high specificity as candidates. Then candidate genes with medium to low expression in female germline cells and low or absent expression in late-stage germline cells and early embryos should be further screened. The classification of high, medium, and low expression levels can be referenced against the expression profiles and drive performance of genes corresponding to promoters that have been tested and used in drive systems within the same species to establish relative standards. These are likely to be somewhat different between species, but general patterns should hold. Thus, it may be possible to identify optimal levels with more limited data in non-model species using our *Drosophila melanogaster* data as a reference.

For species with only bulk RNA-seq data available, it is recommended to prioritize scRNA-seq of reproductive tissues (such as ovary or testis) to improve the efficiency and accuracy of subsequent screening. If scRNA-seq data cannot be obtained, a traditional approach can be adopted. By comparing the expression profiles of reproductive tissues with those of embryos or other somatic tissues, candidate genes with high expression and high specificity in reproductive tissues but low expression in new embryos can be screened. However, this runs the risk of yielding candidates with low female germline expression and high female ovary expression, which may be particularly incompatible when used for a suppression drive (where target genes are often but not always expressed in the ovary), which is the drive type that most needs an optimal promoter. Alternatively, homologous genes that have demonstrated good drive performance and are conserved in other species can be referenced. However, the success rate of identifying suitable promoters using this method is mixed^20,36^. It is possible, though, that using both of these methods together may yield more suitable candidates, each method potentially covering drawbacks of the other.

In summary, we interpreted the performance of a large number of different gene drive Cas9 promoters from a single-cell transcriptomic perspective. Our findings provide a theoretical foundation for the efficient selection of promoter elements and the prediction of drive performance. Specifically, we find that early pre-meiotic germline cells are likely the optimal window for male expression and that the quantitative window for optimal drive performance may be narrow. Avoiding somatic expression may be more complicated and could involve genomic locus. Future work can apply our promoter screening strategy to other organisms, such as disease-vector mosquitoes, in order to optimize drive performance. Additionally, ongoing studies should continue to evaluate the generalizability and applicability of this approach across diverse species and gene drive systems.

## Supporting information

Supplemental Data Sets

## Acknowledgements

This study was supported by the Center for Life Sciences and the National Natural Science Foundation of China (grant 32270672).

## Supplemental Information

**Table S1.** List of donor plasmid names with Cas9 regulatory features.

| Plasmid name | Promoter/5'UTR | Length (bp) | 3'UTR/terminator | Length (bp) |
| --- | --- | --- | --- | --- |
| ISc9ROnG | *rhino* (native site) | N/A | *rhino* (native site) | N/A |
| SNc9ActG | *Act5C* | 2482/1837 | *Act5C* | 1571/121 |
| SNc9banG | *babos* | 440/360 | *nanos* | 839/78 |
| SNc9Cyp6nG | *Cyp6a19* | 1825/14 | *nanos* | 839/78 |
| SNc9fsM3nG | *fs(1)M3* | 2020/90 | *nanos* | 839/78 |
| SNc9fsNnG | *fs(1)N* | 2000/61 | *nanos* | 839/78 |
| SNc9ms1nG | *CG42854* | 444/49 | *nanos* | 839/78 |
| SNc9mtnG | *mtsh* | 419/232 | *nanos* | 839/78 |
| SNc9pp1nG | *CG15262* | 96/139 | *nanos* | 839/78 |
| SNc9pp2nG | *CG15599* | 569/68 | *nanos* | 839/78 |
| SNc9pp3nG | *RpL22-like* | 1896/170 | *nanos* | 839/78 |
| SNc9pp4nG | *CG13577* | 848/87 | *nanos* | 839/78 |
| SNc9pp5nG | *ATPsynεL* | 354/253 | *nanos* | 839/78 |
| SNc9ps2nG | *CG14357* | 452* | *nanos* | 839/78 |
| SNc9ps3nG | *SkpE* | 1915/81 | *nanos* | 839/78 |
| SNc9RO1nG | *rhino* | 371/40 | *nanos* | 839/78 |
| SNc9RO2nG | *rhino* | 1333/40 | *nanos* | 839/78 |
| SNc9RO3G | *rhino* | 3382/40 | *rhino* | 273/100 |
| SNc9s02nG | *CG12477* | 534/145 | *nanos* | 839/78 |
| SNc9sp0nG | *CG6324* | 690/1851 | *nanos* | 839/78 |
| SNc9sp1nG | *CG1894* | 1094/95 | *nanos* | 839/78 |
| SNc9sp2nG | *CG7213* | 1338/124 | *nanos* | 839/78 |
| SNc9sp3_2nG | *CG9692* | 970/44 | *nanos* | 839/78 |
| SNc9sp3nG | *CG11929* | 1570/147 | *nanos* | 839/78 |
| SNc9sp4nG | *CG43106* | 2005/86 | *nanos* | 839/78 |
| SNc9sp5nG | *Prosβ2R2* | 128/92 | *nanos* | 839/78 |
| SNc9spLnG | *CG32814* | 64/1062 | *nanos* | 839/78 |
| SNc9tejnG | *tej* | 897/108 | *nanos* | 839/78 |
| SNc9UbiG | *Ubi-p63E* | 204/1425 | *Ubi-p63E* | 209/102 |
| SNc9320nG | *CG32068* | 1036/31 | *nanos* | 839/78 |

Naming convention: SN = split nuclease, c9 = Cas9, G = EGFP
For length of Promoter/5' UTR, the first number represent the number of nucleotides before the 5' UTR, and the second represents the number of nucleotides in the 5' UTR. For length of the 3' UTR/terminator, the first number represents the length of the 3' UTR, and the second represents the number of nucleotides included after the 3' UTR.
\*Due to lack of detailed genomic annotation, represents the total 5' UTR/promoter length.

**Table S2.**
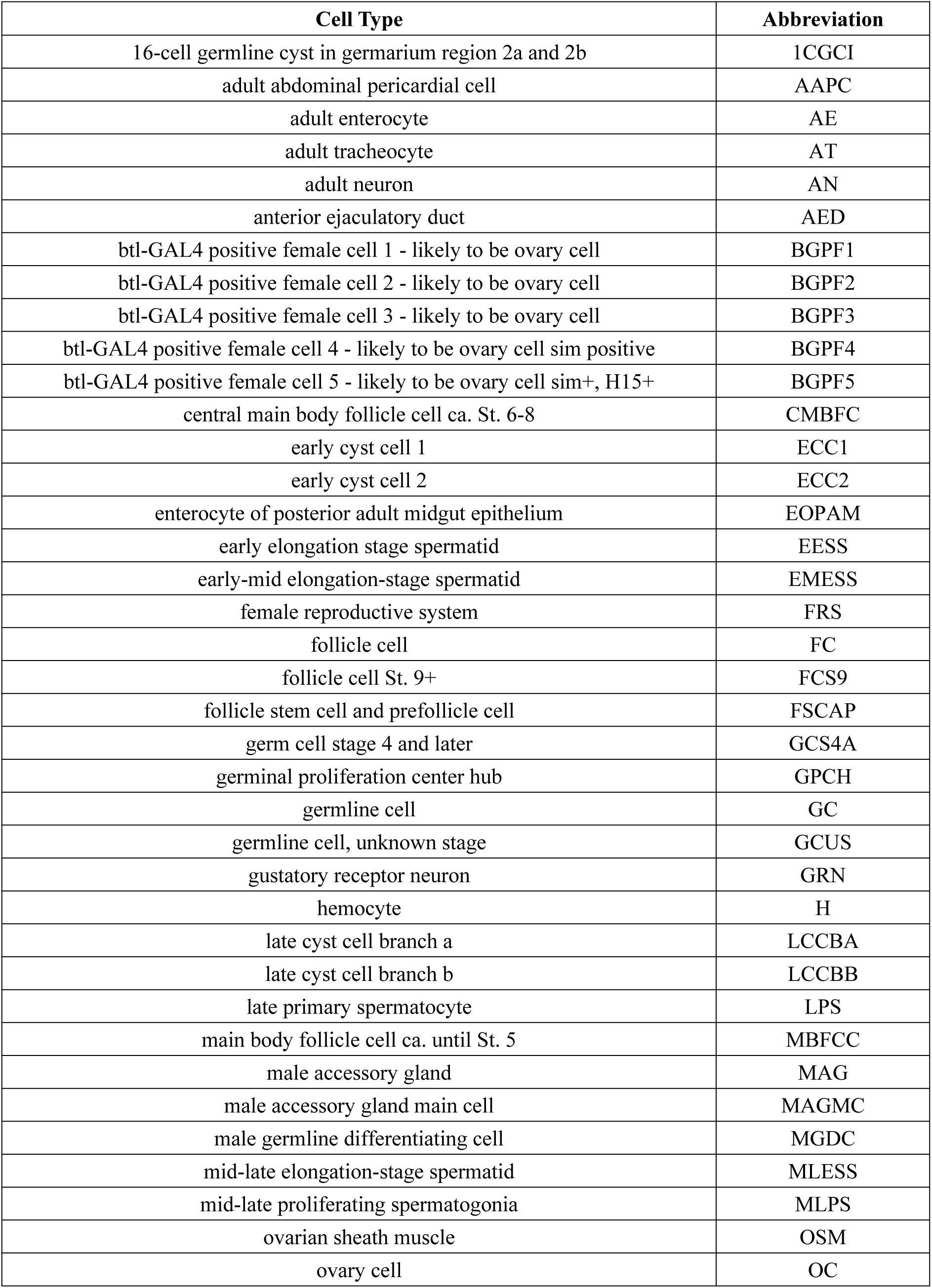
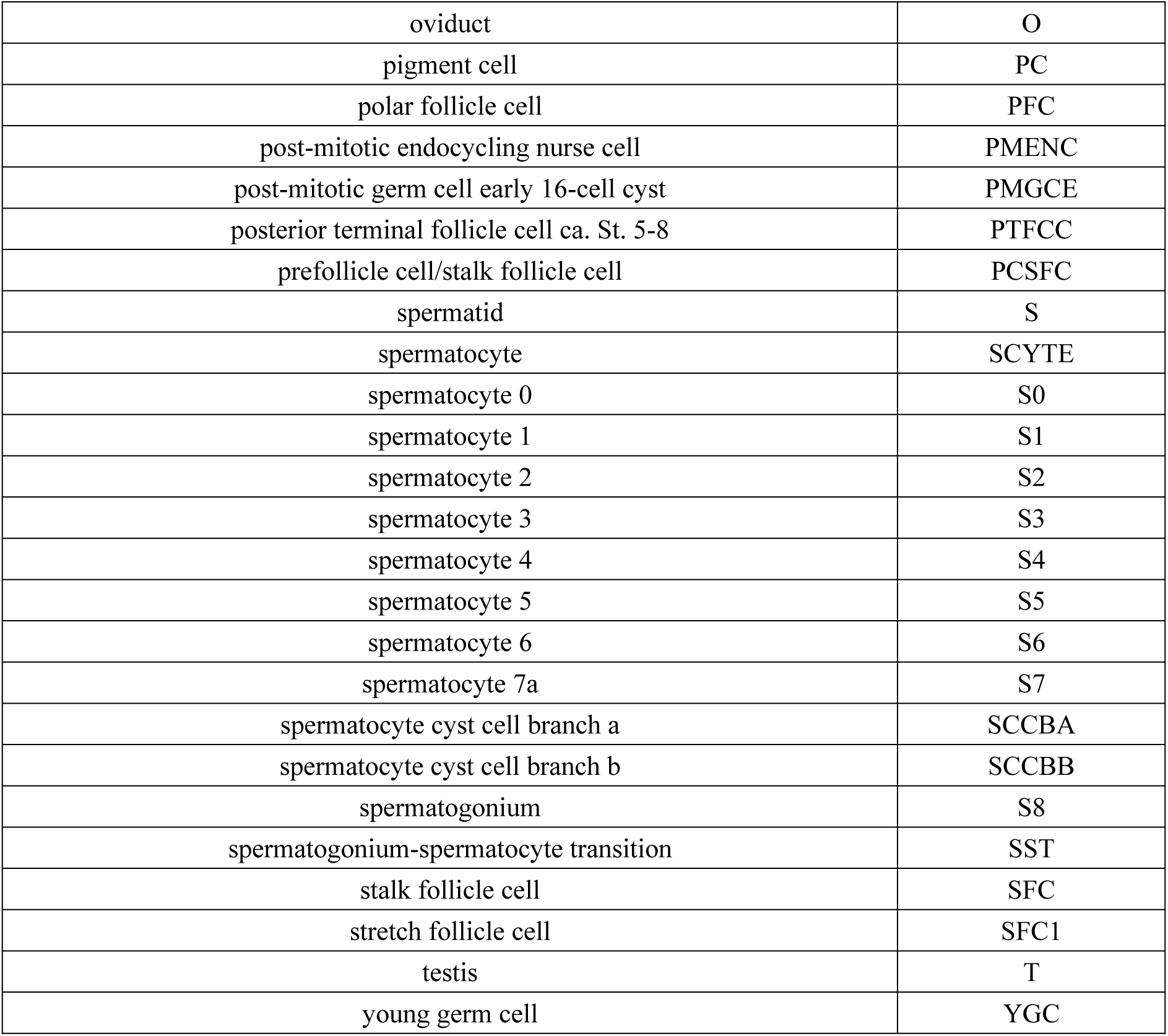
Cell Type Abbreviations.

**Figure S1.**
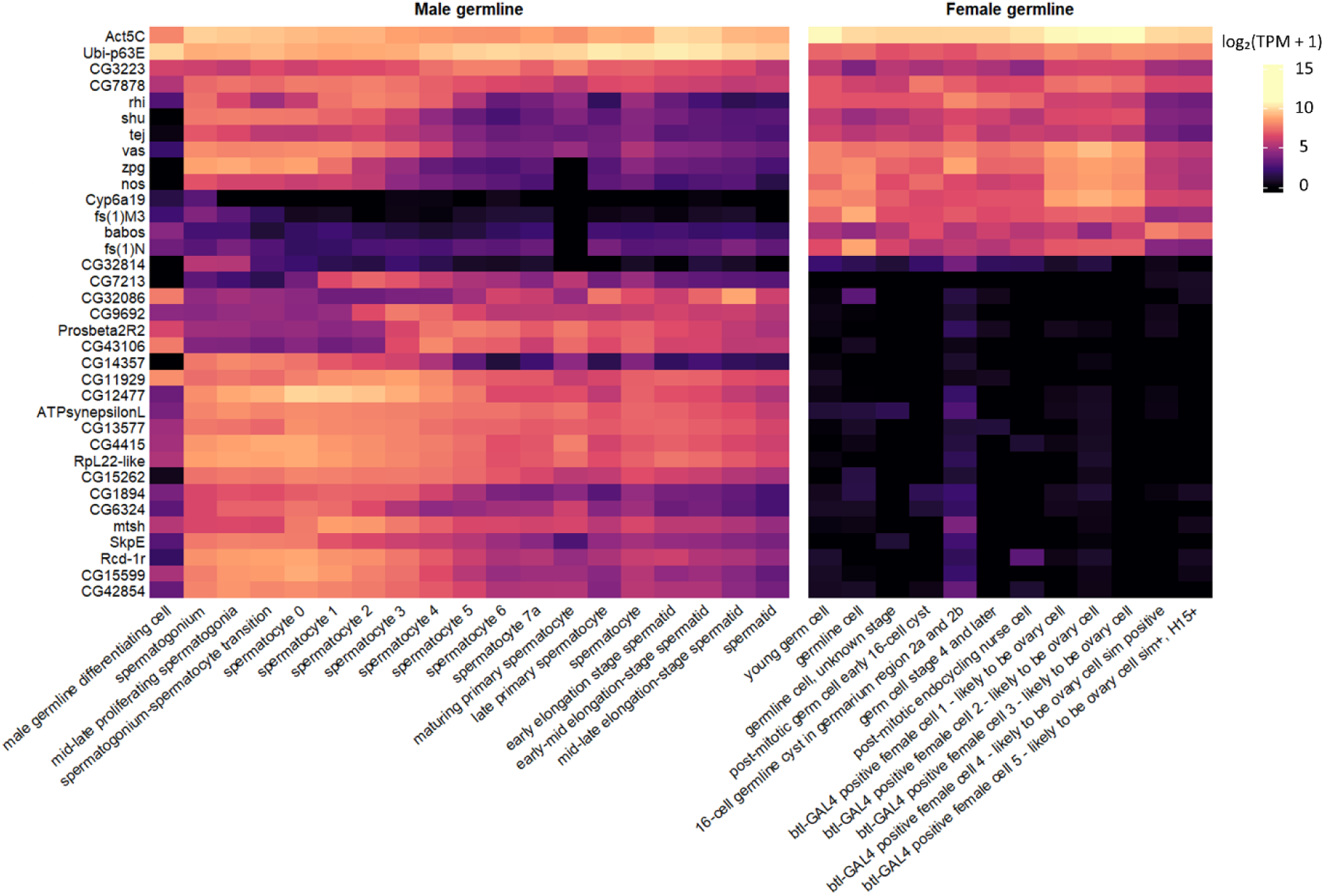
Germline cell expression profiles of promoter-candidate genes in *Drosophila melanogaster*. The heatmap displays the expression profiles of 35 gene candidates across 259 single cell subtypes from the Fly Cell Atlas. Raw Transcripts Per Million (TPM) were transformed to log_2_(TPM + 1) and color-mapped. To determine gene order, Euclidean distances were computed complete-linkage hierarchical clustering. Cell types were reordered to best approximate temporal position in gamete development (earlier cells are to the left).

**Figure S2.**
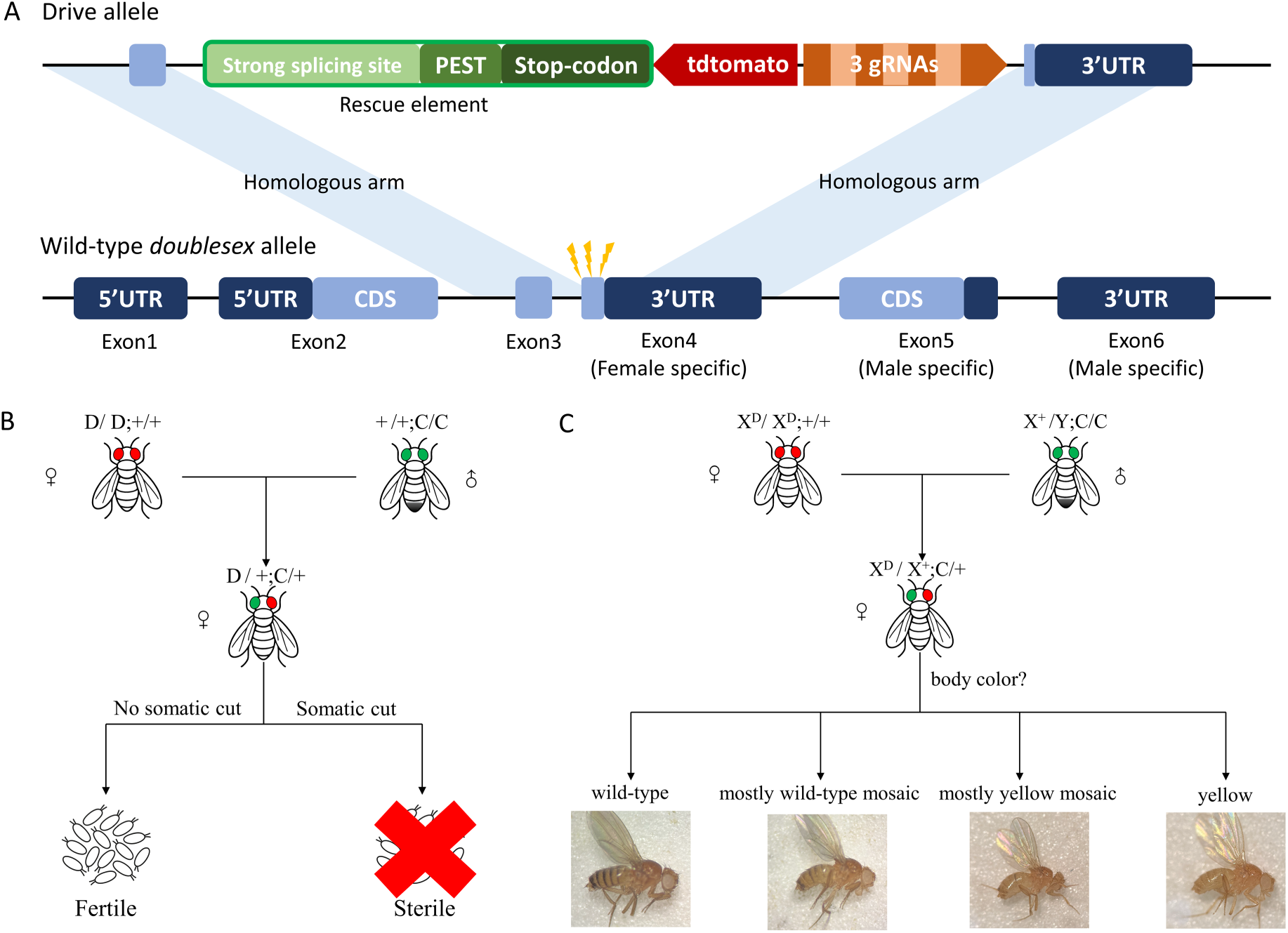
Assessment of somatic expression in the *doublesex* and *yellow* systems. **A.** The *doublesex* system includes two components: a split Cas9 element and the drive element. The split Cas9 element is as previously described. The drive element contains three gRNAs targeting female-specific exon 4 *doublesex*, which are linked by tRNAs and expressed by the U6:3 promoter. Females with disrupted *doublesex* in both chromosomes have an intersex phenotype. **B.** The cross scheme of *doublesex* drive system for evaluating somatic expression. Females with the drive element are first crossed with males with the Cas9 element. Females carrying both elements are evaluated by crossing them with wild-type males and assessing their fertility (none of these females displayed increased intersex characteristics). **C.** The cross scheme used to evaluate somatic expression in the *yellow* drive system was identical to that used for the *doublesex* system. In the *yellow* system, somatic expression was evaluated by the body color of females carrying both fluorescent markers. More yellow pigmentation indicated higher somatic expression. Related data are provided in Data Set S2.

**Figure S3.**
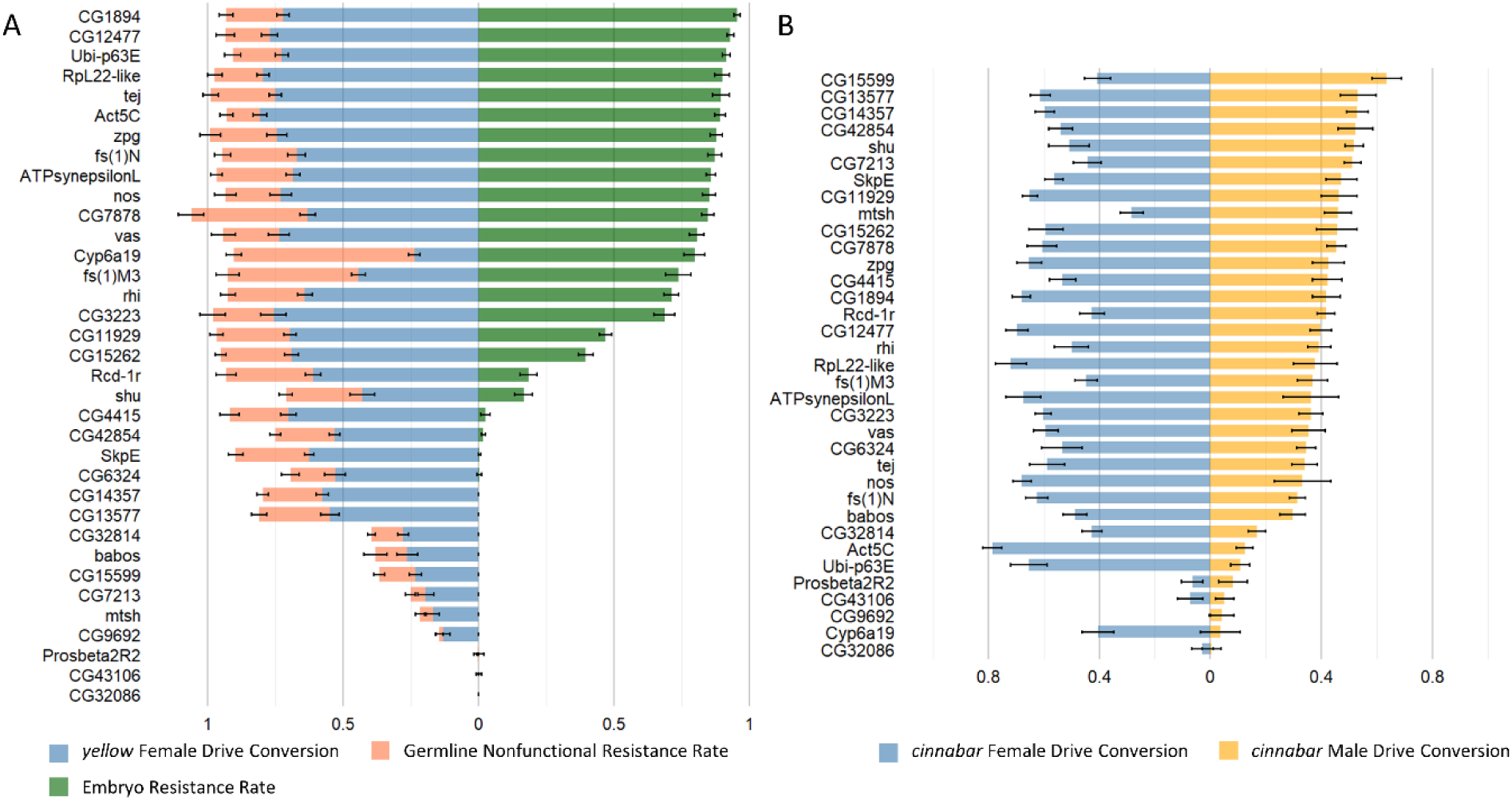
Drive performance of Cas9 constructs with different promoters in the *yellow* and *cinnabar* system. **A.** Drive performance of 35 Cas9 constructs regulated by different gene promoters in the *yellow* system. The sum of the drive conversion rate and the germline nonfunctional resistance rate is nearly equal to the total germline cut rate for each promoter. For visualization purposes, calculated drive conversion rates below zero are displayed as zero. Source data are provided in Data Sets S1-S2. **B.** Drive performance of 35 Cas9 constructs regulated by different gene promoters in the *cinnabar* system. Error bars indicate standard error of the mean. Source data are provided in Data Set S3.

**Figure S4.**
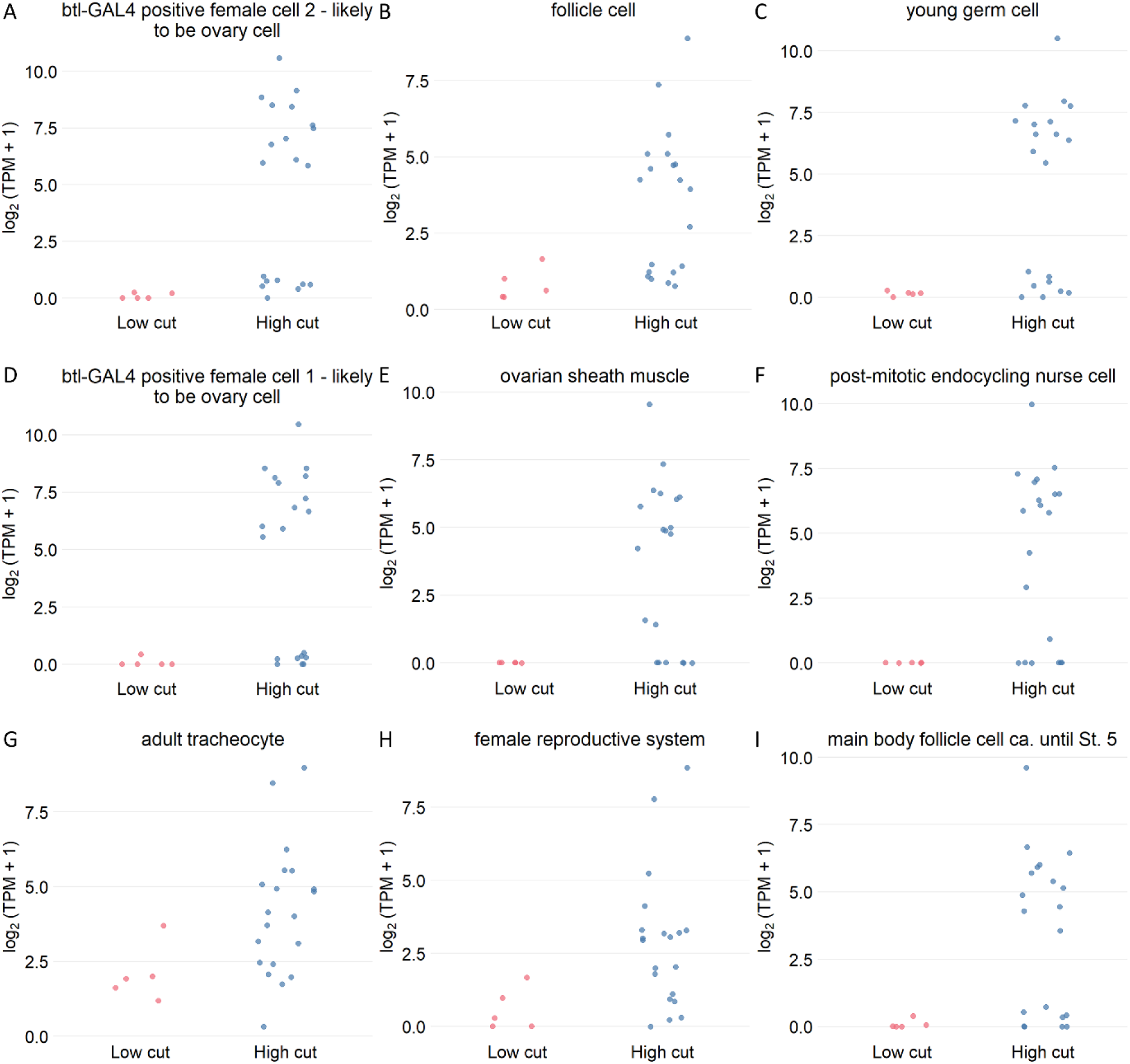
Expression differences of promoter-associated genes between high and low total germline cut rate groups in the *yellow* system. Scatter plots showing the log_2_(TPM + 1) expression of promoter-associated genes in significantly different cell types (*p* < 0.05), comparing high cut rate (greater than 90%) and low cut rate (less than 25%) groups. Statistical differences were evaluated using the Wilcoxon rank-sum test. Only the top nine most statistically significant cell types are shown. Each point represents the expression of one gene in the cell type. The vertical axis shows the log_2_(TPM + 1) expression of a promoter’s gene.

**Figure S5.**
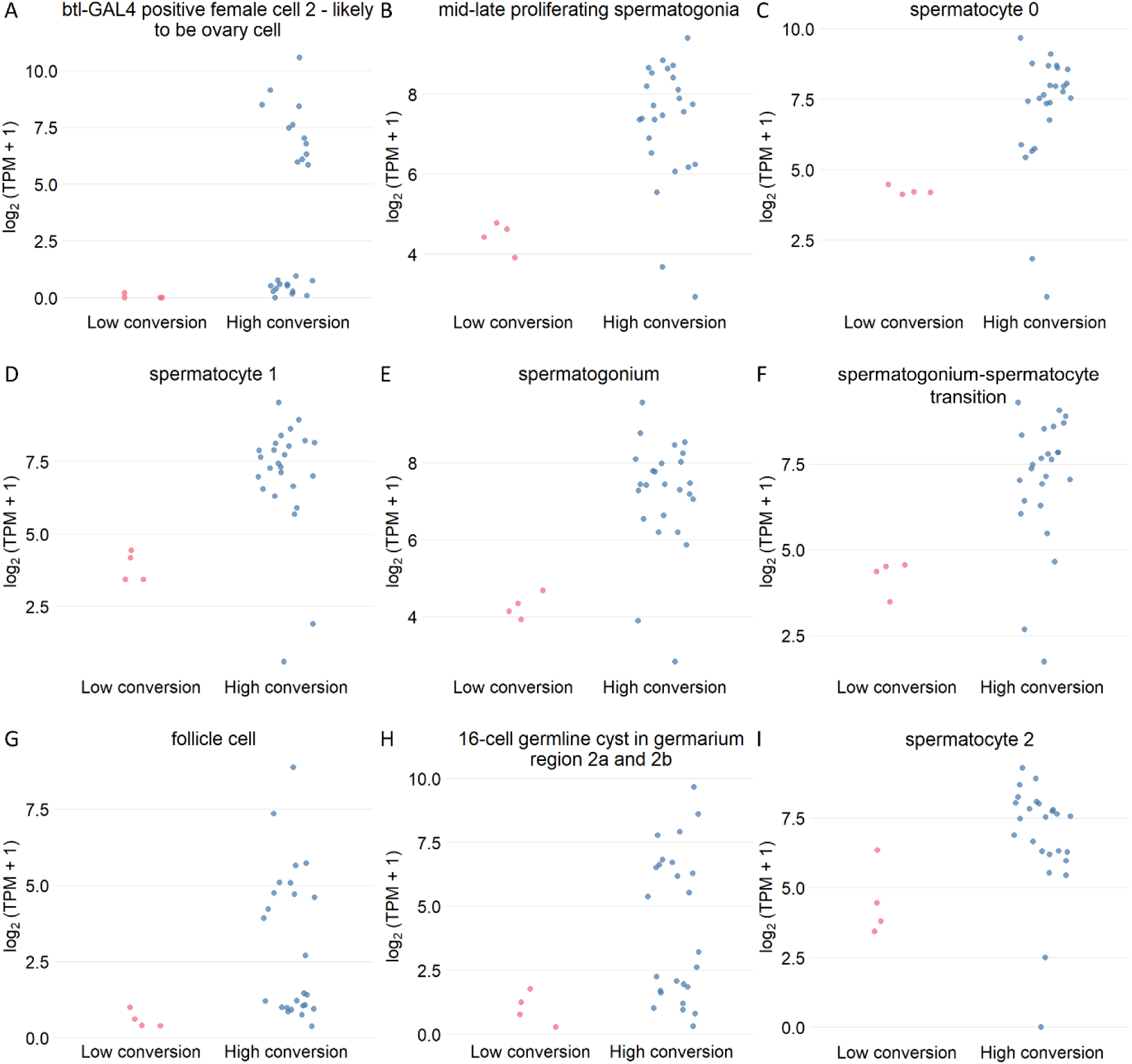
Expression differences of promoter-associated genes between high and low drive conversion rate groups in the *yellow* system. Scatter plots showing the log_2_(TPM + 1) expression of promoter-associated genes in significantly different cell types (*p* < 0.05), comparing high drive conversion (over 40%) and low drive conversion (under 15%) groups. Statistical differences were evaluated using the Wilcoxon rank-sum test. Only the top nine most statistically significant cell types are shown. Each point represents the expression of one gene in the cell type. The vertical axis shows the log_2_(TPM + 1) expression of a promoter’s gene.

**Figure S6.**
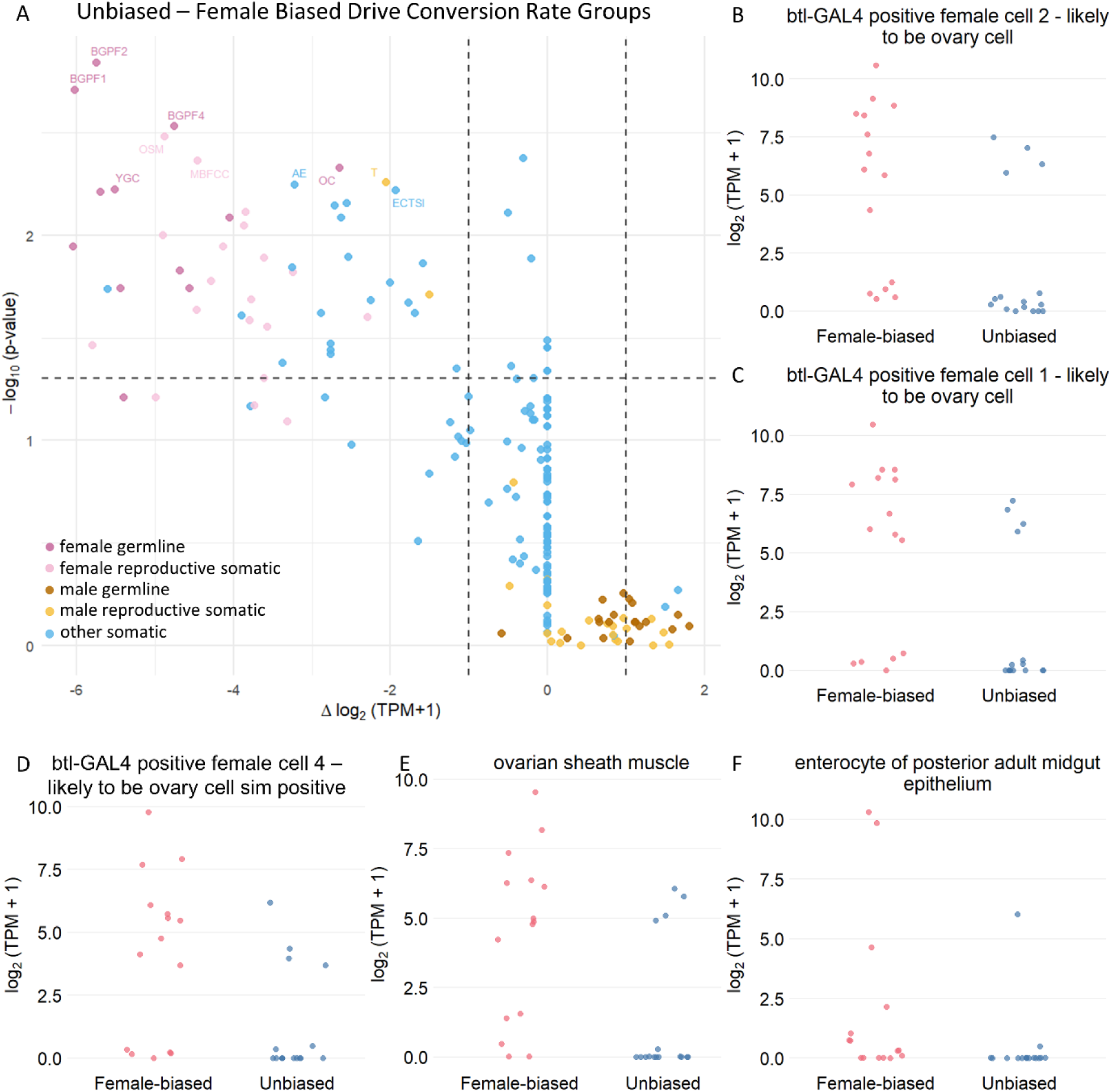
Female-biased conversion is associated with distinct promoter-associated gene expression across cell types. **A.** Differential expression analysis of promoter associated genes between female-biased and unbiased groups across various cell types. Each point represents a single cell type from the Fly Cell Atlas dataset, with the horizontal axis indicating the difference in log_2_(TPM+1) between unbiased and female-biased groups, and the vertical axis showing the -log_10_(*p*-value) derived from Wilcoxon rank-sum tests. Several cell types with significant differences (*p* < 0.05 and |Δlog_2_(TPM+1)| > 1) are labeled with their abbreviated names (Table S2). Source data are provided in Data Set S9. **B-F.** Expression differences of promoter-associated genes between female-biased and unbiased groups. Scatter plots showing the log_2_(TPM + 1) expression of promoter-associated genes in significantly different cell types (*p* < 0.05), comparing female-biased (female drive conversion rate significantly exceeds that of males) and unbiased (female drive and male drive conversion rate do not differ significantly) groups as measured in the *cinnabar* system.

**Figure S7.**
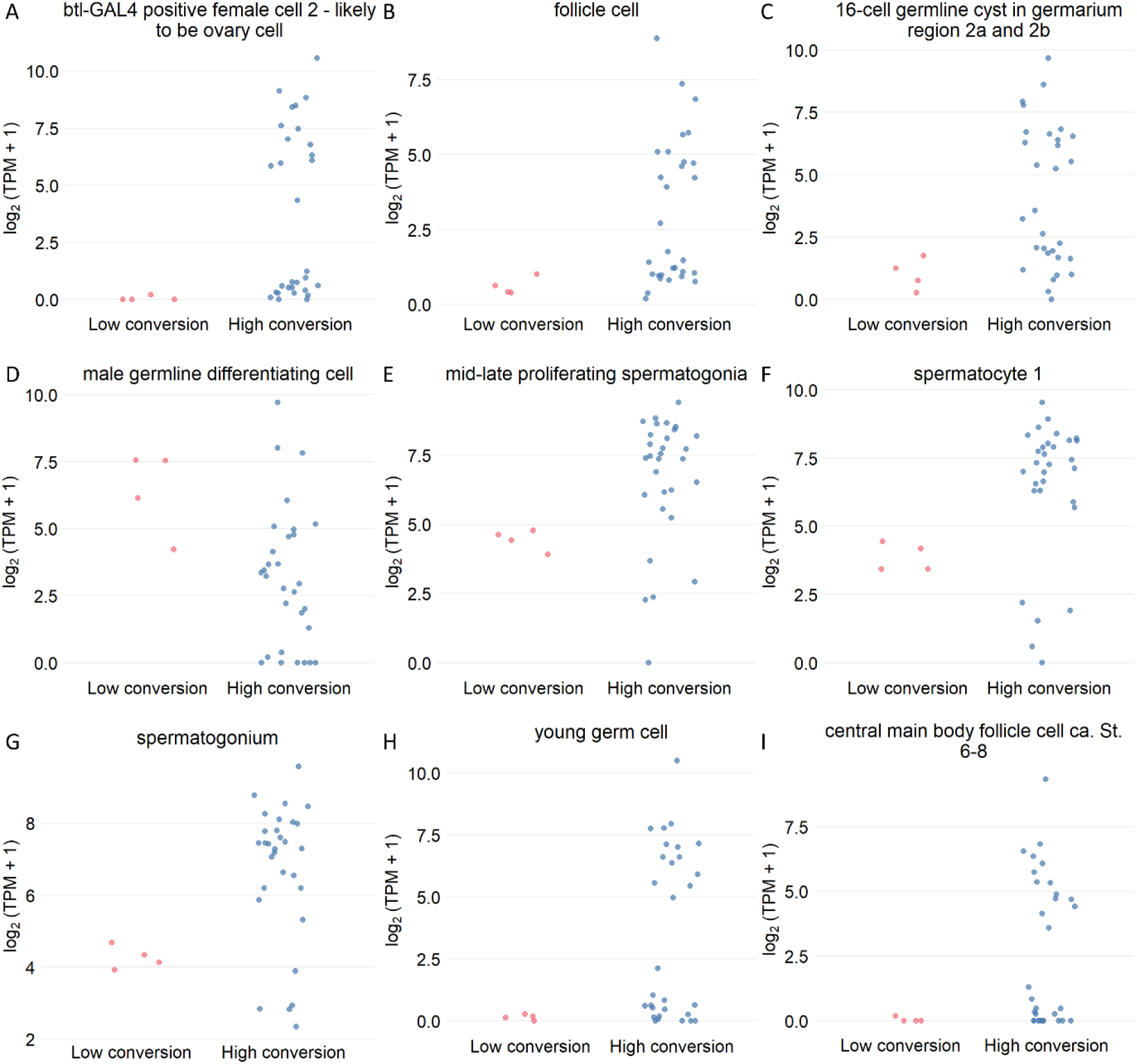
Expression differences of promoter-associated genes between high and low female drive conversion rate groups in the *cinnabar* system. Scatter plots showing the log_2_(TPM + 1) expression of promoter-associated genes in significantly different cell types (*p* < 0.05), comparing high drive conversion (greater than 40%) and low drive conversion (less than 15%) groups. Statistical differences were evaluated using the Wilcoxon rank-sum test. Only the top nine most statistically significant cell types are shown. Each point represents the expression of one gene in the cell type. The vertical axis shows the log_2_(TPM + 1) expression of a promoter’s gene.

**Figure S8.**
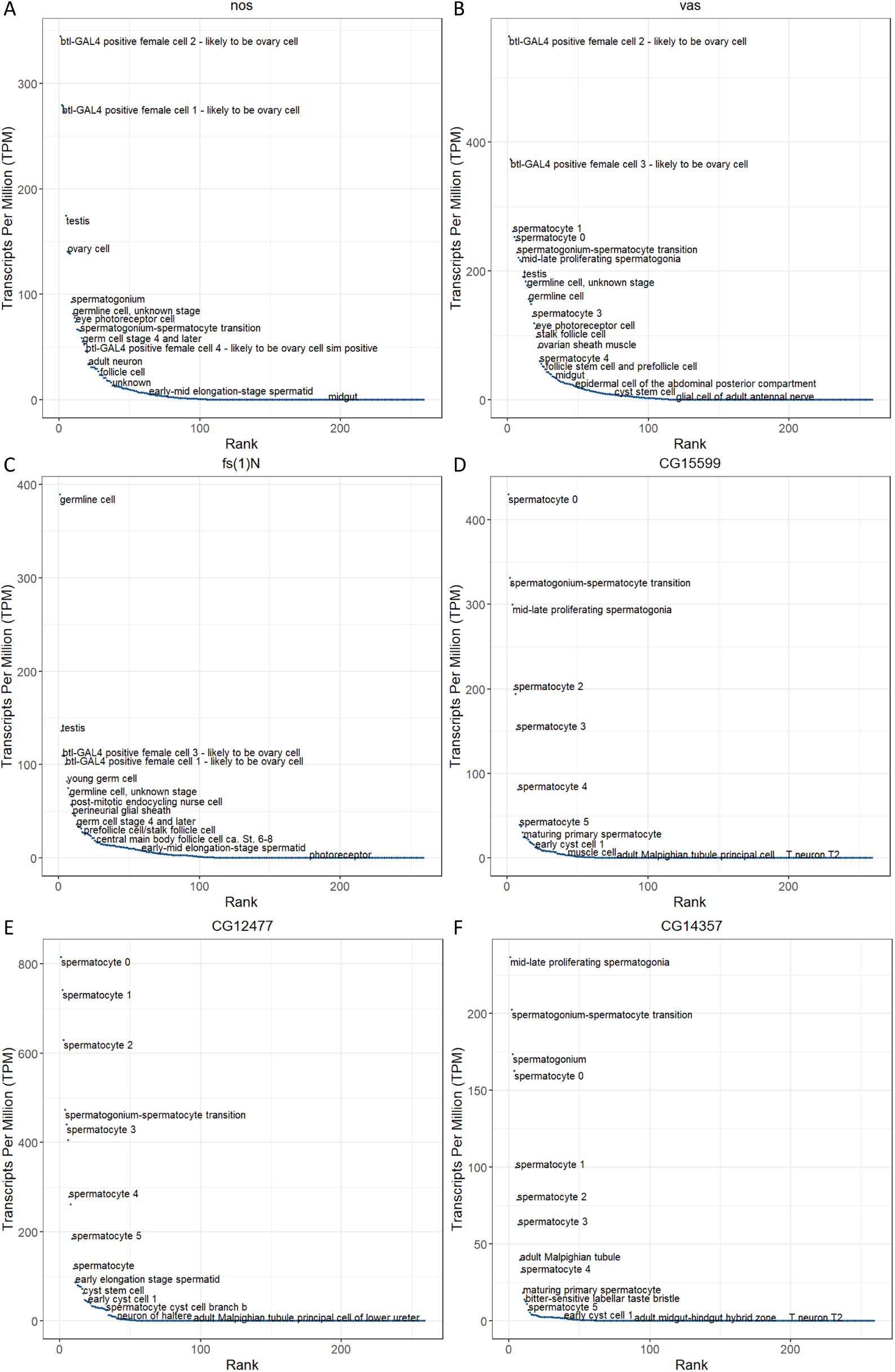
Ranked expression distribution of different genes across single cells. Each point represents the Transcripts Per Million of different genes in a single cell type, ranked in descending order.

**Figure S9.**
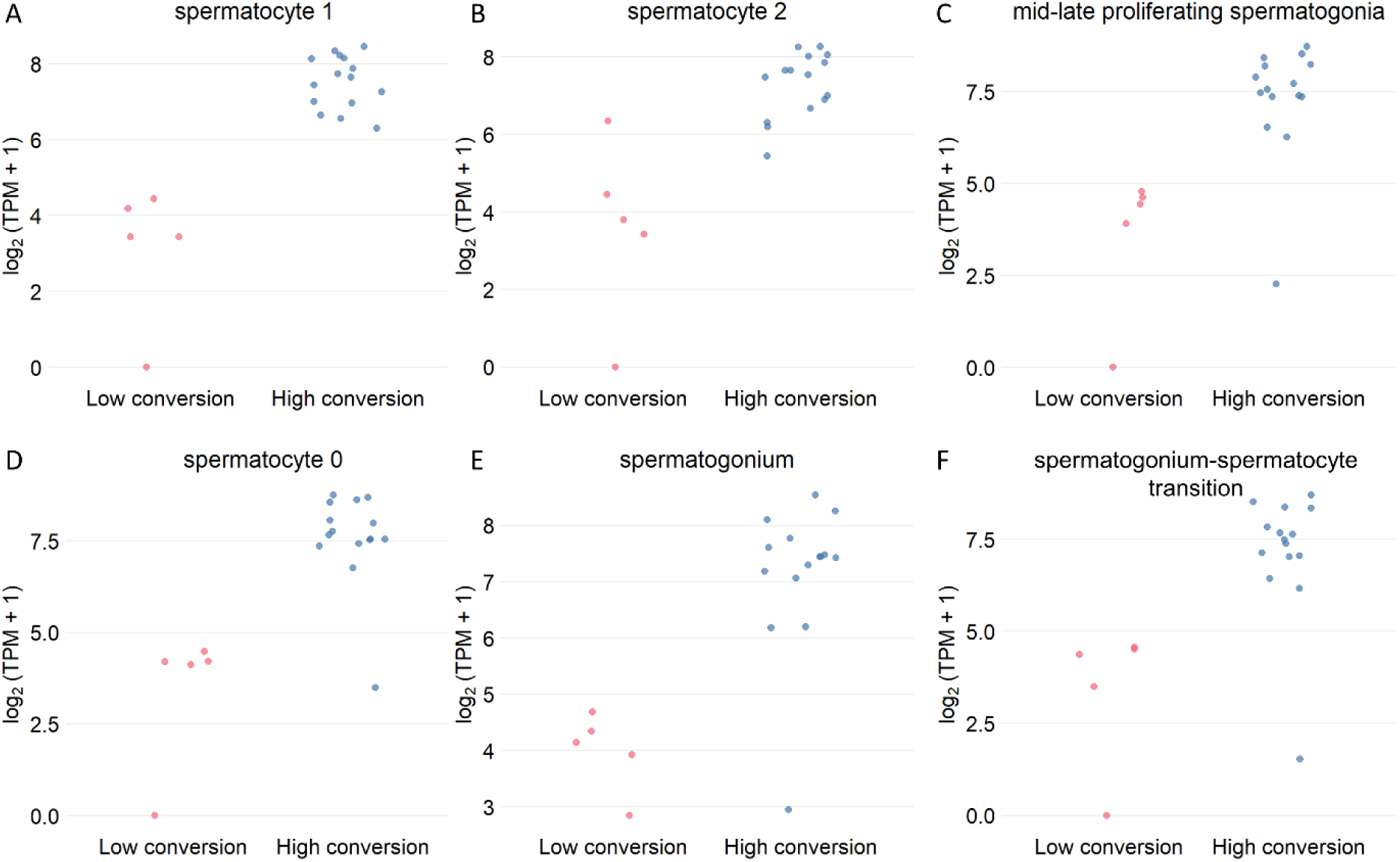
Expression differences of promoter-associated genes between high and low male drive conversion rate groups in the *cinnabar* system. Scatter plots showing the log_2_(TPM + 1) expression of promoter-associated genes in significantly different cell types (*p* < 0.05), comparing high conversion (over 40%) and low conversion (under 10%) groups. Statistical differences were evaluated using the Wilcoxon rank-sum test. Only the top nine most statistically significant cell types are shown. Each point represents the expression of one gene in the cell type. The vertical axis shows the log_2_(TPM + 1) expression of a promoter’s gene.

**Figure S10.**
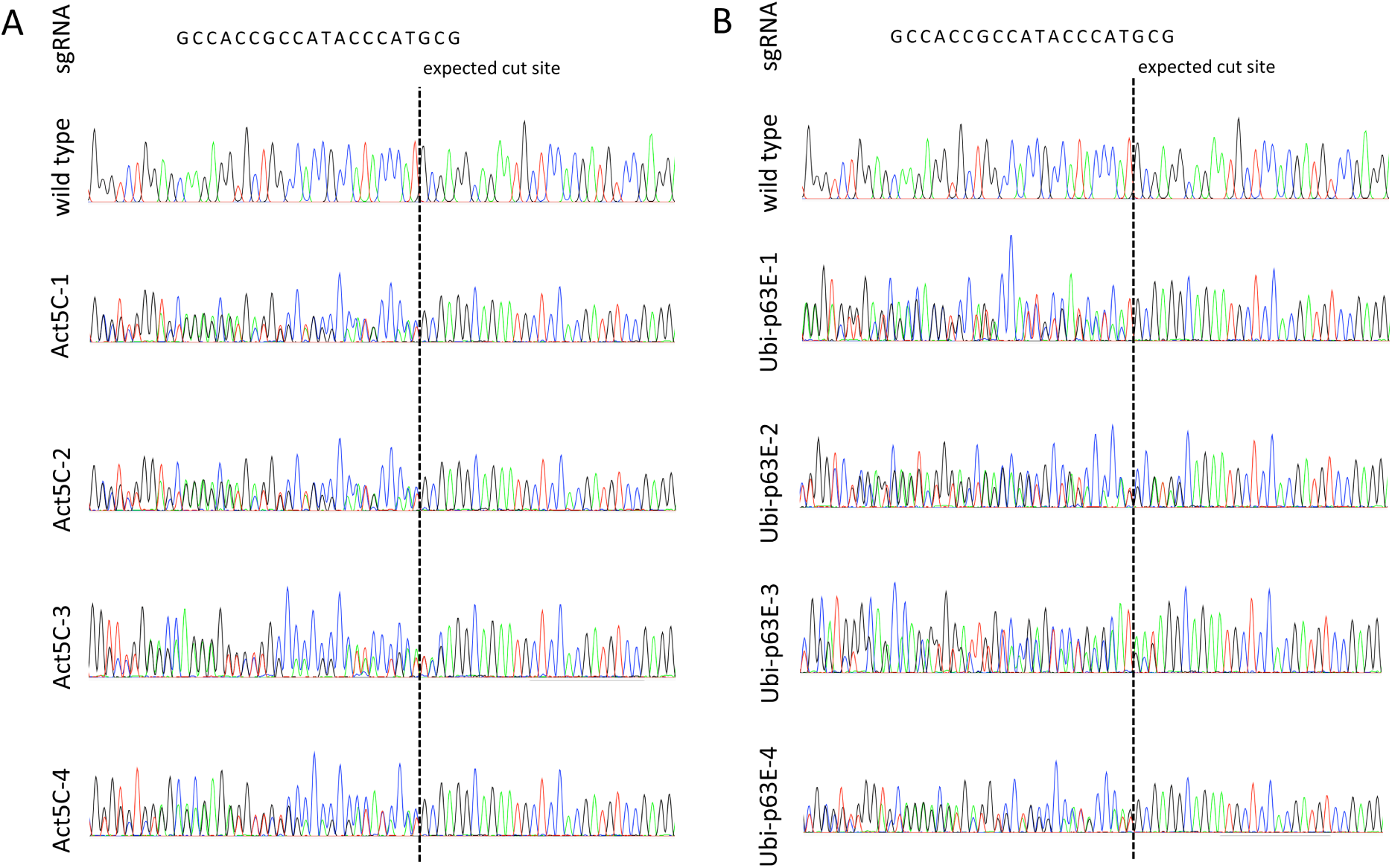
Target site sequencing at the *cinnabar* locus. **A.** Target site sequencing in non-drive progeny from male parents with both the *Act5C* Cas9 element and *cinnabar* drive element. **B.** Target site sequencing in non-drive progeny from male parents with both the *Ubi-p63E* Cas9 element and *cinnabar* drive element. The dashed line indicates the expected Cas9 cut site. Mixed sequencing peaks or heterozygous signals at the target region (present in all four tested of each group) suggest that cleavage events occurred, forming a germline resistance allele in the parent that was inherited by the sequenced offspring.

**Figure S11.**
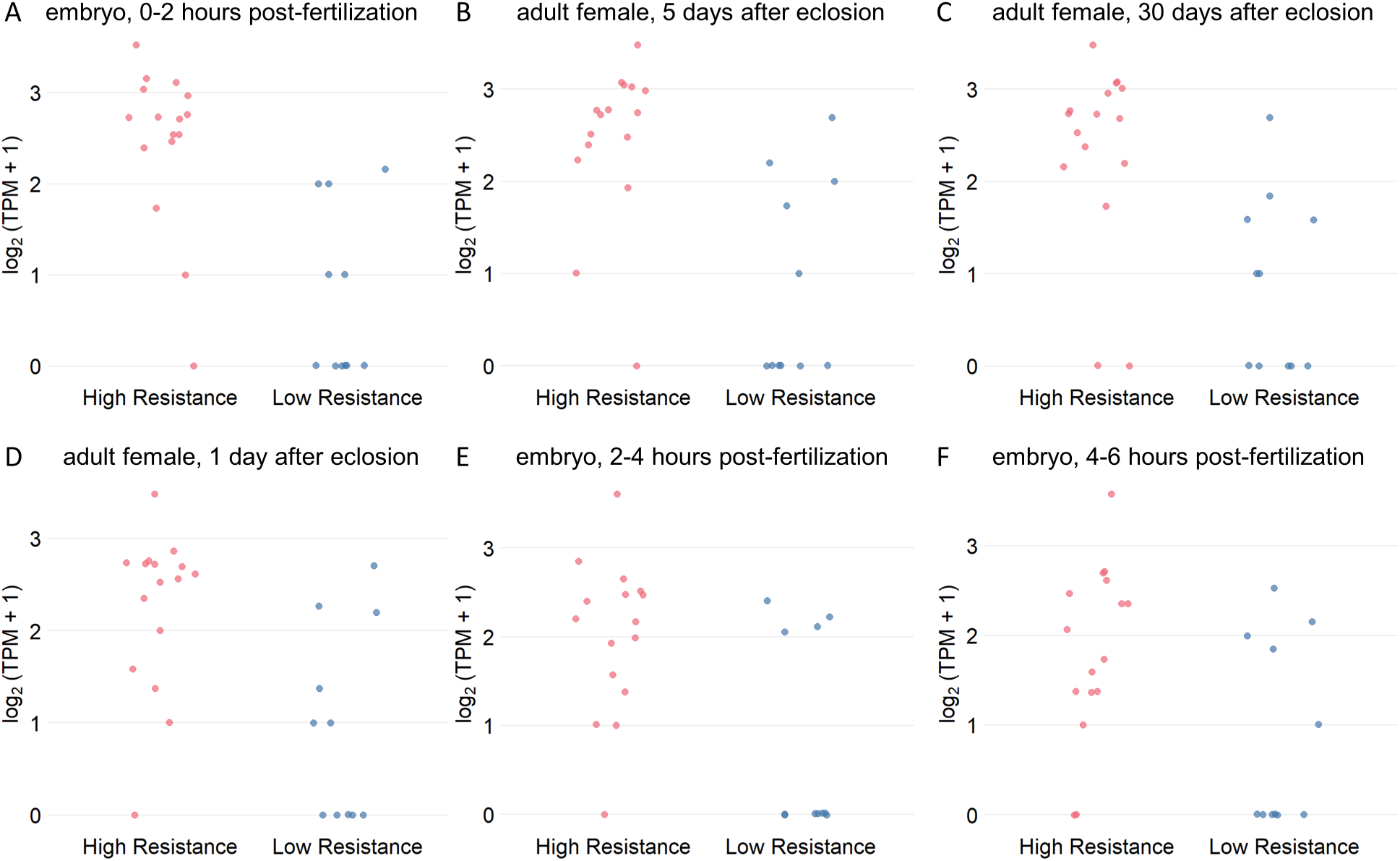
Expression differences of promoter-associated genes between high and low embryo resistance rate groups in the *yellow* system, based on bulk RNA-Seq data. All statistically significant cell types are shown. Each point represents the expression of one promoter-associated gene in the sample. Plot structure and interpretation as in Fig.S2. Source data are provided in Data Set S12.

**Figure S12.**
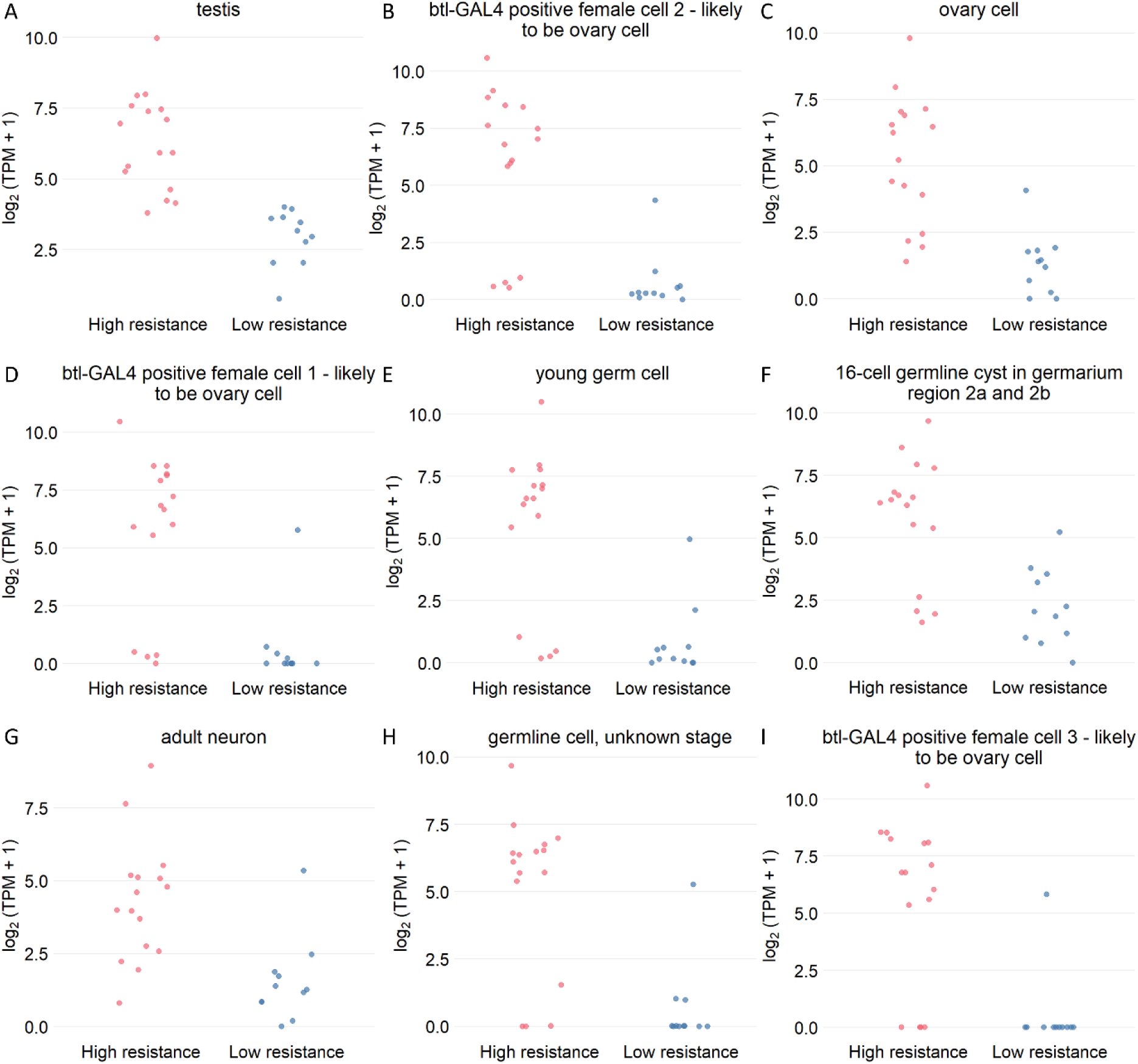
Expression differences of promoter-associated genes between high and low embryo resistance rate groups in the *yellow* system, based on single-cell transcriptomic data. Scatter plots showing the log_2_(TPM + 1) expression of promoter-associated genes in significantly different cell types (*p* < 0.05), comparing high embryo resistance (greater than 65%) and low embryo resistance rate (less than 5%) groups. Only constructs with drive conversion rates greater than 40% were included. Statistical differences were evaluated using the Wilcoxon rank-sum test. Only the top nine most statistically significant cell types are shown. Each point represents the expression of one gene in the cell type. The vertical axis shows the log_2_(TPM + 1) expression of a promoter’s gene.

**Figure S13.**
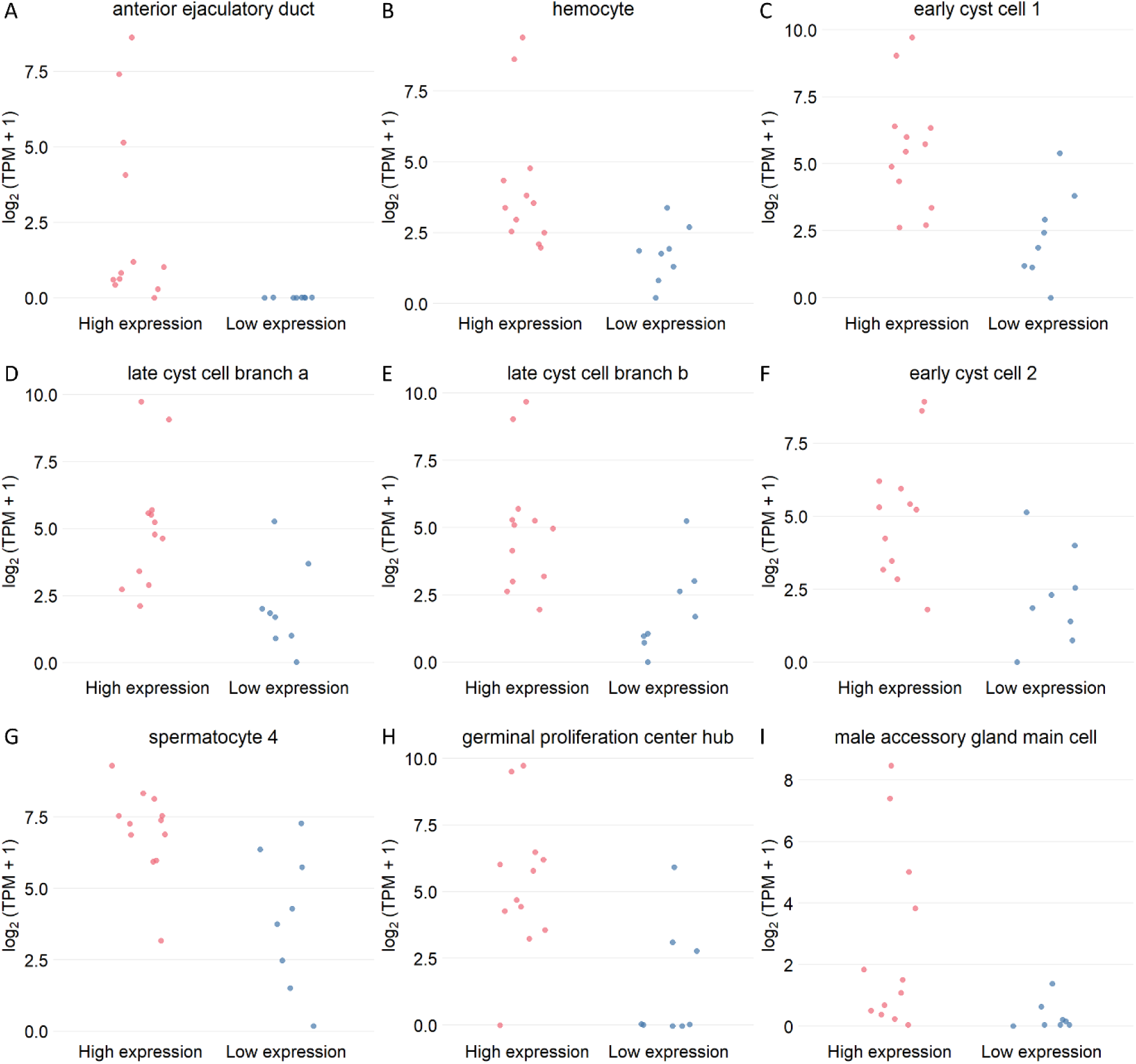
Expression differences of promoter-associated genes between high and low somatic expression groups in the *yellow* system. Scatter plots showing the log_2_(TPM + 1) expression of promoter-associated genes in significantly different cell types (*p* < 0.05), comparing low somatic expression (wild-type) and high somatic expression (yellow phenotype). Only constructs with drive conversion rates greater than 40% were included. Statistical differences were evaluated using the Wilcoxon rank-sum test. Only the top nine most statistically significant cell types are shown. Each point represents the expression of one gene in the cell type. The vertical axis shows the log_2_(TPM + 1) expression of a promoter’s gene.

**Figure S14.**
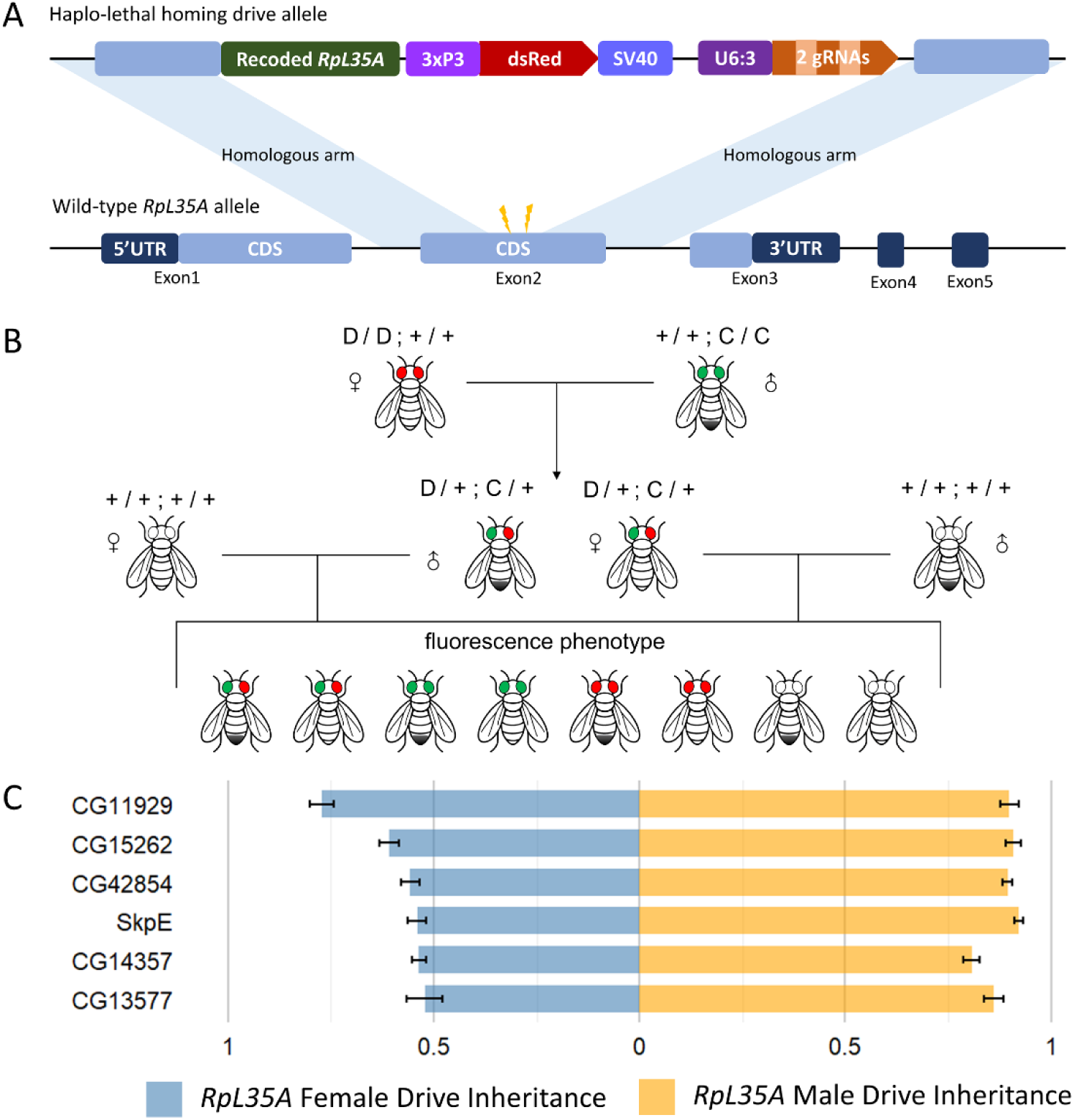
The split drive system targeting *RpL35A.* **A.** The *RpL35A* system includes two components: a split Cas9 element and the drive element.. The drive element contains two gRNAs targeting exon 2 of *RpL35A*, which are linked in tandem via a tRNA and expressed by the U6:3 promoter. *RpL35A* is a haplolethal gene encoding a highly conserved protein component of the 60S ribosomal subunit, so nonfunctional resistance alleles result in nonviability, regardless of the other allele. **B.** The cross scheme of the *RpL35A* drive system. Females with the drive element strain are first crossed to males with the Cas9 elements. Female or males carrying both fluorescent markers are selected from the progeny and crossed to *w^1118^* flies. Drive performance was assessed by analyzing the fluorescence phenotypes of the resulting offspring. **C.** Drive performance of six Cas9 constructs regulated by different gene promoters in the *RpL35A* system. Error bars indicate standard error of the mean. Source data are provided in Data Set S4.

## Notes

https://github.com/jchamper/Single-Cell-Promoters-Homing

### Competing Interest Statement

The authors have declared no competing interest.

